# TurboID-based proximity-dependent labeling using SOBIR1 as a bait in potato leads to the identification of novel defense-related signaling partners

**DOI:** 10.64898/2026.07.19.739134

**Authors:** Tatiana Marti Ferrando, Sergio Landeo Villanueva, Sjef Boeren, Matthieu H.A.J. Joosten, Vivianne G.A.A. Vleeshouwers

## Abstract

The plant immune system comprises a complex signaling network that is activated upon perceiving molecules derived from invading pathogens. The first line of defense at the plant cell surface is mediated by receptor-like proteins (RLPs) and receptor-like kinases (RLKs). RLPs, which lack a cytoplasmic signalling domain themselves, constitutively interact with the RLK SUPRESSOR OF BIR1-1 (SOBIR1), which is a key component initiating immune signal transduction upon pathogen perception. Therefore, elucidating the composition of the SOBIR1 protein complex will contribute to understanding the basic molecular mechanisms of plant disease resistance. Most of the studies focused on the identification of SOBIR1-interacting proteins are limited to model plants, due to technical challenges and lack of reliable genome and proteome databases in crop plants. Here, we evaluate the application of the biotin ligase TurboID (TbID)-based proximity-dependent labeling (PL) approach by transiently expressing SOBIR1 from *Nicotiana benthamiana* (NbSOBIR1), fused to TbID in leaves of the wild potato *Solanum microdontum*. We show that NbSOBIR1-YFP-TbID properly accumulates in potato and that proximal proteins are biotinylated. Quantitative proteomic analysis yielded over 130 candidate proteins to be in the proximity of the cytoplasmic kinase domain of NbSOBIR1, of which some could be linked to disease resistance by KEGG pathway and gene ontology (GO) molecular function analysis. We also studied the dynamics of the proteome in proximity of NbSOBIR1 upon perception of the INF1 elicitin of *Phytophthora infestans* that was co-expressed in potato with the elicitin receptor ELR, which is an RLP that constitutively interacts with SOBIR1. We found more than 80 proteins, including the NB-LRR REQUIRED FOR HR-ASSOCIATED CELL DEATH 1 (NRC1), putatively interacting with NbSOBIR1. In conclusion, we were able to successfully apply PL in potato and a future roadmap for further research on deciphering the composition of protein complexes involved in immune signaling has been established.

## Introduction

Most studies on plant immunity have been approached in a reductionist manner [1]. The immune receptor network of plants consists of a diverse array of receptors that recognize pathogen-derived molecules and often function in a coordinated manner, with crosstalk and interdependency between different components of the network [2, 3]. This complex and highly interconnected system has multiple layers of regulation and signaling pathways to adapt to different pathogen challenges. The first layer of defense is activated upon recognition of apoplastic effectors or conserved microbe- or pathogen-associated molecule patterns (MAMPs/PAMPs) by cell-surface receptors such as pattern recognition receptors (PRRs) and leads to pattern-triggered immunity (PTI) [4, 5]. The second layer of defense, which is a more specific and robust response, is activated upon recognition of cytoplasmic effectors by resistance (R) proteins, leading to the activation of effector-triggered immunity (ETI) [4].

PRRs comprise receptor-like kinases (RLKs) and receptor-like proteins (RLPs) that reside at the plant cell surface and function together with co-receptors as a receptor [6]. PRRs contain an extracellular domain, which in most cases consists of leucine-rich repeats (LRRs) that sense external proteinaceous signals from the pathogen. The ectodomain is followed by a transmembrane (TM) domain and a short C-terminal tail in case of RLPs, and a cytoplasmic kinase domain in case of RLKs. The kinase domain has a signaling function, as its phosphorylation activates the defense signaling cascade [7]. RLPs form a bimolecular RLK by constitutively interacting with the RLK SUPPRESSOR OF BIR1-1 (SOBIR1) and initiate downstream signalling through the kinase domain of SOBIR1 [8, 9]. SOBIR1 is conserved in all plant species and plays a key role in immunity by regulating the activity of RLPs and their signaling partners [8]. The RLP/SOBIR1 pair, as well as RLKs, interact with the BRASSINOSTEROID INSENSITIVE1 (BRI1)-ASSOCIATED KINASE1 (BAK1), also known as SOMATIC EMBRYOGENESIS RECEPTOR KINASE3 (SERK3) upon ligand perception [10–13]. BAK1/SERK3 activates the PRR complex and phosphorylates receptor-like cytoplasmic kinases (RLCKs), which play a role in activating downstream signaling pathways [14, 15]. Once active, RLCKs like BOTRYTIS-INDUCED KINASE1 (BIK1), phosphorylate signaling proteins, leading to the activation of defense responses such as the release of reactive oxygen species (ROS) produced by the RESPIRATORY BURST OXIDASE HOMOLOGUE (RBOH) and the activation of MITOGEN-ACTIVATED PROTEIN KINASE (MAPK) cascades [16, 17].

Potato late blight is one of the most devastating plant diseases and is caused by the hemi-biotrophic oomycete pathogen *Phytophthora infestans*. This pathogen is highly adaptable and rapidly evolves new strains that are resistant to current crop protection strategies [18, 19]. The introgression of *R* genes encoding cytoplasmic nucleotide-binding LRR proteins (NLRs) derived from wild potato species has been widely applied over the last years, although this approach does not effectively provide long-term resistance to late blight yet [20]. Being at the frontline of defense, PRRs are another source to be exploited for more durable and broad-spectrum resistance. The RLP ELICITIN RECEPTOR (ELR) was the first PRR cloned from the wild potato species *Solanum microdontum*, which has been reported as an important source of late blight resistance [21–27]. Elicitins are a family of secreted proteins that are structurally conserved among *Phytophthora* and *Pythium* species and that play a role in sequestering sterols and other lipids from plant membranes. Their structure holds a hydrophobic pocket with the ability to bind sterols, which is essential for growth and sporulation of these sterol auxotrophic pathogens. Elicitins share features with MAMPs and they can elicit a hypersensitive response (HR) in *Nicotiana* and various *Solanum* species upon their recognition [28], yet they modulate plant immune responses independently of their sterol carrier capacity [28, 29]. *P. infestans* encodes seven elicitins (INF1, INF2A, INF2B, INF3, INF4, INF5 and INF6), with INF1 being the most highly produced elicitin *in vitro* [30, 31]. ELR recognizes four of these elicitins, namely INF1, INF2A, INF5 and INF6, and this broad-spectrum recognition renders ELR a potential candidate to reach more durable resistance to *P. infestans*.

Plant immunity is based on a complex system with multiple layers of regulation and signaling pathways responding to pathogen challenges. The signaling network relies on swift interactions between different proteins to ensure the proper cellular responses, which in a specific context will culminate in programmed cell death, referred to as the HR [32]. The study of protein-protein interactions (PPIs) *in planta* should offer potential targets for molecular breeding. Methods such as yeast two-hybrid (Y2H) and co-immunoprecipitation (co-IP), have their limitations as being laborious, generating a high false-positive rate, and having difficulties with detecting weak or short-lived interactions. Proximity-dependent labeling (PL) has been successfully developed to study PPIs in animals, micro-organisms and in the model plants *Arabidopsis* and *Nicotiana benthamiana* [33–36]. PL experiments allow the identification of proteins that are in close proximity to a bait protein of interest, and a recently engineered biotin ligase, referred to as TurboID (TbID), has been described as a suitable candidate for PL in plants. TbID has been applied to investigate the profile of the subcellular proxitome and signaling pathways triggered by immune receptors in *Arabidopsis* and *N. benthamiana* [37–43], transcription factor interactions in *Solanum lycopersicum* (tomato) hairy roots [44], and effector-host protein interactions in *Zea mays* (maize) [40]. Thus, the TbID-based PL tool has been established in plant science and will be very useful to understand the function and regulation of protein complexes in different biological contexts.

In this study we applied TbID-based PL, followed by mass spectrometry (MS) to identify the biotinylated proteins, using transient expression of NbSOBIR1-YFP-TbID as a bait in leaves of potato plants. PL-MS upon agro-co-infiltration of ELR and INF1, together with NbSOBIR1-YFP-TbID in potato leaves, was also performed. Several candidate NbSOBIR1 interactors were found, and we speculate on the potential involvement of some of them in plant immunity. Nevertheless, more research needs to be conducted with complementary studies, such as virus-induced gene silencing (VIGS), gene editing or other complementary PPI methods such as co-IP or Y2H to confirm which of the candidates are real SOBIR1 interactors and are truly involved in plant immunity. With our studies we have added information supporting our understanding of the composition of the SOBIR1-signaling network in mounting resistance in potato, that might provide novel targets for future molecular breeding for late blight resistance.

## Results

### Establishment of TbID-based proximity labeling in a wild *Solanum* genotype

To implement TbID-based PL by the *Agrobacterium*-mediated transient expression system, we first identified a suitable *Solanum* genotype to be employed for this. According to our experience, agroinfiltration in potato leaves might result in specific resistance responses triggered by the protein that is expressed, but might also cause nonspecific defense responses to *Agrobacterium* [45]. We screened 25 progeny genotypes of the F1 population 3521, which has *Solanum microdontum* (MCD360-1) and *Solanum verrucosum* in its pedigree [24]. Many of the tested wild *Solanum* progeny genotypes showed chlorosis or necrosis upon agro-infiltration of the empty vector (EV)-pK7WG2 (negative control), or failed to show specific cell death upon agro-co-infiltration of ELR-pK7WG2 and INF1-pK7WG2 (positive control), and we considered them not amenable for agroinfiltration (**Table S1-S2**). However, the wild *Solanum* hybrid 3521-321 (MCD-321) did not show visible symptoms upon agro-infiltration of EV-pK7WG2 and INF1-pK7WG2, whereas agro-co-infiltration of ELR-pK7WG2 and INF1-pK7WG2 resulted in a confluent cell death at 5 days post infiltration (dpi) (**Table S1, Figure 1A**). Therefore, we identified the hybrid MCD-321 to be well amenable to our PL experiments to study proximal protein interaction networks with TbID-fused NbSOBIR1 (A0A0H3U2A6) as a bait.

**Figure 1.**
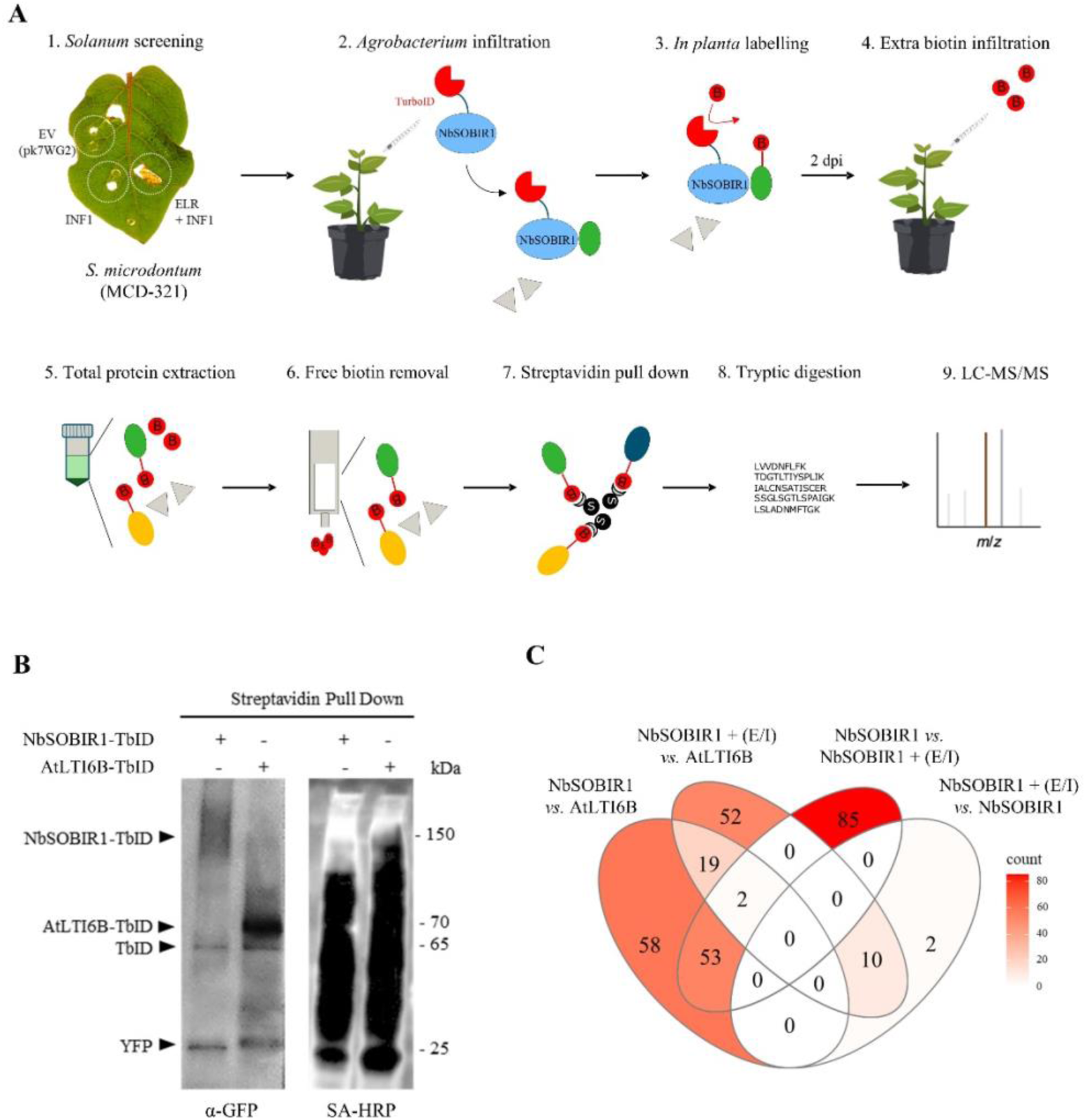
Roadmap for the identification of the proxitome of NbSOBIR1 in *S. microdontum*, using a TurboID-based approach. **A. Experimental design.** (1) *Solanum* screening to identify a compatible genotype for *Agrobacterium* transient expression using co-agroinfiltration of ELR and INF1 as a positive control; and infiltration of INF1 and the empty vector (EV) pK7WG2 as a negative control. (2) *Agrobacterium* transient expression of NbSOBIR1 fused to the TurboID enzyme (NbSOBIR1-TurboID). (3) The proxitome of NbSOBIR1-TurboID is biotinylated by TurboID, using the endogenous biotin from the plant. (4) Addition of 200 μM biotin at 48 hours after agro-infiltration to increase the biotinylation of proteins. (5) Collect samples 1 hour after biotin infiltration and isolation of total proteins using RIPA-based buffer. (6) Free biotin removal from the samples with de-salting columns to improve the efficiency of the streptavidin pull down. (7) Capture of the biotinylated proteins with streptavidin beads. (8) Tryptic digestion of the captured proteins. (9) Separation of the generated peptides by liquid chromatography (LC) and identification of the proteins by mass spectrometry (MS). **B. Streptavidin pull down.** Proteins fused to TurboID are accumulating in potato MCD-321 (shown using ⍺-GFP antibodies) and are able to biotinylate proximal proteins *in planta* (shown using Streptavidin (SA)-HRP protein). **C. Number of significantly detected proteins proximal to NbSOBIR1, in different comparisons.** Three treatments were performed: expression of NbSOBIR1-YFP-TbID (NbSOBIR1), expression of NbSOBIR1-YFP-TbID combined with ELR and INF1 (NbSOBIR1 + (E/I)) and the control AtLTI6B-YFP-TbID (AtLTI6B). Each treatment consisted of three independent replicates. The Venn diagram represents the numbers of putatively interacting proteins with the protein of interest of all possible comparisons of the different treatments.

To set up a pipeline for identifying the proximal proteome of the NbSOBIR1-containing receptor complex in the wild potato hybrid MCD-321, we transiently expressed NbSOBIR1 C-terminally fused to the yellow fluorescent protein (YFP) and TbID. As a reference for non-specific biotinylation and plasma membrane (PM) localization, we used the *Arabidopsis thaliana* LOW TEMPERATURE INDUCED PROTEIN 6B (AtLTI6B), also C-terminally fused to YFP-TbID (**Figure S1)**. AtLTI6B is a small membrane-associated protein (6 kDa), and it has been previously used as a PM marker in other studies [46, 47]. We first proved that proper accumulation of both TbID-fused proteins took place in leaves of MCD-321 by agroinfiltration. At 2 dpi, a 200 μM biotin solution was infiltrated in the same leaves and 1 hour later the leaves were harvested. After total protein extraction, samples were desalted to remove the free biotin present in the extracts and biotinylated proteins were subsequently pulled down with streptavidin (SA) beads (**Figure 1A**). Subsequent analysis by western blotting (WB) with GFP antibodies confirmed the expression of NbSOBIR1-YFP-TbID, which presents a smear at around 150 kDa, likely indicative of protein complex formation or uncontrolled cross-linking of proteins with this co-receptor (**Figure 1B**) [48]. The AtLTI6B-YFP-TbID (70 kDa) control fusion protein also accumulated successfully in the wild potato MCD-321 upon its agro-infiltration (**Figure 1B**). Bands corresponding to free TbID-YFP (65 kDa) and YFP (27 kDa) were also detected on the WB. Additionally, using Streptavidin (SA) protein conjugated to horseradish peroxidase (HRP) enzyme (SA-HRP) revealed a broad smear of biotinylated proteins in both samples, confirming *in planta* catalytic activity of the TbID-fusion protein (**Figure 1B**).

To validate the capture of the bait protein by mass spectrometry (MS), we analyzed the tryptic peptides originating from the NbSOBIR1-YFP-TbID pull-downs. We identified 42 peptides matching NbSOBIR1, which mapped to both the LRR and kinase domains (data not shown). Resulting peptides from the transmembrane (TM) domain are highly hydrophobic and difficult to detect by LC-MS [49]. To account for potential sequence overlap with endogenous potato orthologues, we performed a BLAST search against the *S. tuberosum* proteome (UniProt ID: UP000011115), thereby identifying two orthologues: M1CKT4 (StSOBIR1; 82.2% identity) and M1B7X0 (StSOBIR1-like; 76.5% identity). While most peptides were shared due to high sequence similarity, we identified unique peptides specific to the endogenous StSOBIR1 and StSOBIR1-like proteins (14 peptides each). These results indicate that the PL assay enriches both the recombinant bait and potentially its endogenous potato orthologues, likely due to the formation of constitutive homodimers, as previously reported [7]. MS analysis also confirmed the presence of YFP and TbID in the respective samples (**Table S1**). However, AtLTI6B was not detected, likely because the size of the peptides is below the detection limit or due to the highly hydrophobic nature of the resulting peptides. Altogether, we demonstrated that NbSOBIR1-YFP-TbID and the control AtLTI6B-YFP-TbID proteins are functional and accumulate in leaves of MCD-321 upon their transient expression, validating the suitability of the system for proteomic profiling.

### Identification of NbSOBIR1-interacting proteins using TbID-based PL

To identify putative interactors of NbSOBIR1, we performed a comparative proxitome analysis of NbSOBIR1-YFP-TbID versus AtLTI6B-YFP-TbID (**Figure 2A**). Statistical analysis (FDR < 0.05) revealed 132 NbSOBIR1-YFP-TbID-enriched putative interactors, when compared to the AtLTI6B control (**Table 1**, **Figure 1C**). The identified proximal proteins were predominantly associated with plant-pathogen interactions and signal transduction, mainly localizing to the PM and cytosol (**Figure 2B-C**). We identified a robust set of signaling components, including 27 RLKs, 5 calcium-dependent protein kinases (CDPKs), and key signal transmission proteins such as 14-3-3s, remorins, and the RPM1-INTERACTING PROTEIN 4 (RIN4). Notably, highly significant interactors (log2-fold change > 1.5 and -log_2_ p-value > 2.5) contained four RLPs, including homologues of the *Cladosporium fulvum* resistance proteins Cf-9 (Hcr9-9D and Hcr9-OR2A), and Cf-2 (Hcr2-p3) [50]. The fourth RLP is a LRRNT_2 domain-containing protein (M1CW41), which is involved in defence responses in plants. Additionally, CDPK1 and proteins involved in the ubiquitin pathway (F-box family proteins and E3 ubiquitin transferases) were grouped among the most differentially abundant proteins. Subcellular localization analysis indicated a predominant enrichment of PM (52%) and cytoplasmic (C) (28%) proteins, consistent with the expected localization of the SOBIR1 complex, although proteins associated within other compartments were also detected (**Table 1**, **Figure 2D**). Nineteen of these proteins are reported to be present in multiple compartments, e.g. the blue light photoreceptors phototropin-1 (M1CDM5) and phototropin-2 (M1C1T9) are located in the PM, C, Golgi (G) and chloroplast (CP) (**Table 1**). In conclusion, the identification of biotinylated proteins in the NbSOBIR1-YFP-TbID-generated sample revealed potential candidate proteins that might be interacting directly or indirectly with NbSOBIR1 and are mainly related to processes playing a role in plant-pathogen interactions.

**Figure 2.**
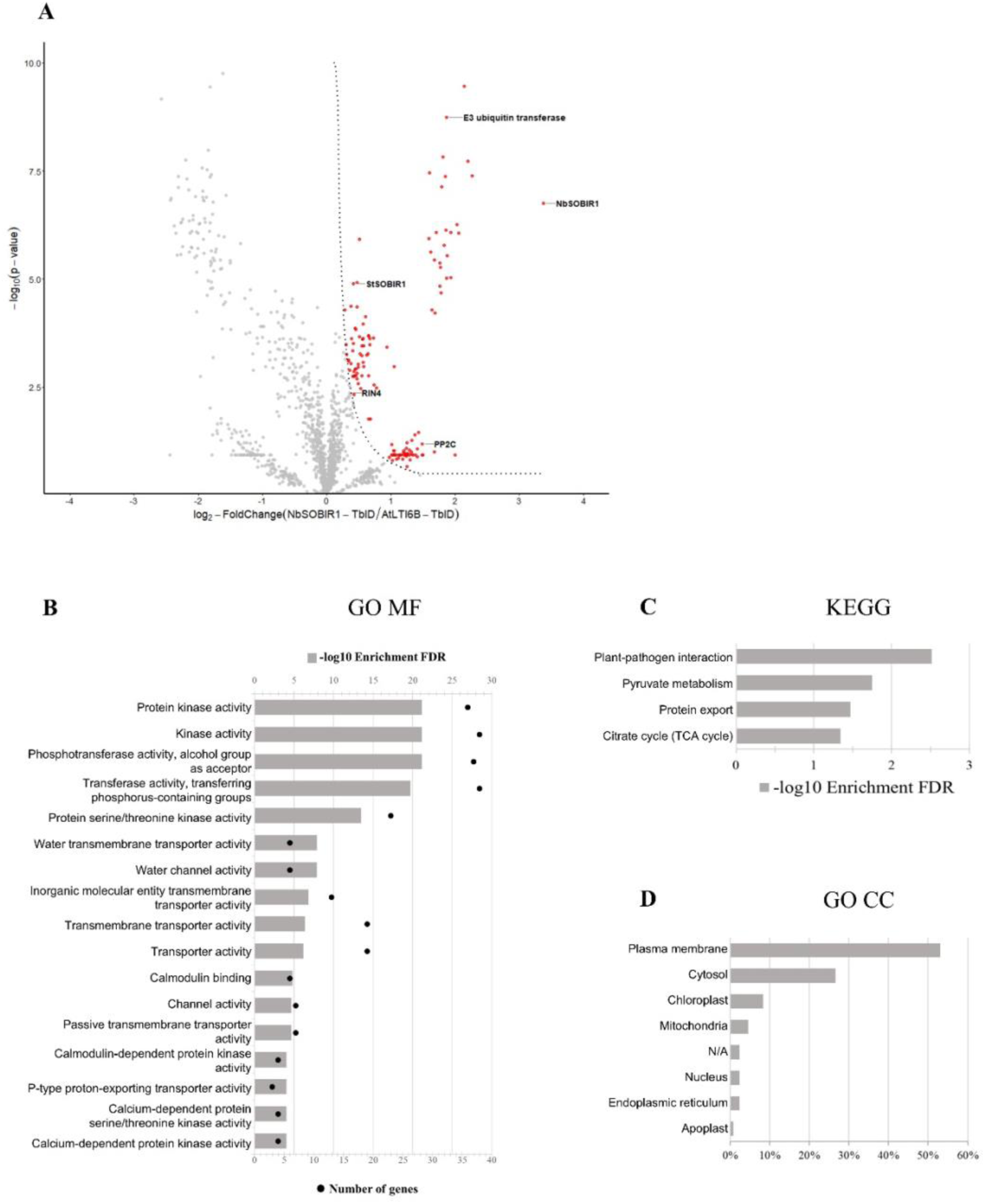
Detecting proteins proximal to NbSOBIR1 in potato MCD321. **A.** The Volcano plot depicts the log2-fold change versus the -log2 p-value, with SOBIR1-TbID on the right and AtLTI6B-YFP-TbID on the left. The fold change is the LFQ value of the identified proteins between SOBIR1-TbID and AtLTI6B-YFP-TbID samples. The p-value is the probability of a protein to be significantly proximal to SOBIR1-TbID or to AtLTI6B-YFP-TbID. Significant proteins biotinylated by NbSOBIR1-YFP-TbID were selected using an S0 of 0.1 and a p-value lower or equal to 0.05 (marked by red dots). The grey dots represent non-significant identified proteins, with an S0 lower than 0.1 and a p-value higher than 0.05. **B.** Gene Ontology (GO) enrichment analysis of Molecular Function (MF) from significant identified proteins upon transient expression of NbSOBIR1-YFP-TbID. Pathways with an FDR lower or equal to 0.05 are represented by -log10 enrichment FDR as a grey bar. The numbers of genes involved in each pathway are represented by black dots. **C.** KEGG pathway enrichment analysis of significant identified proteins upon transient expression of NbSOBIR1-YFP-TbID. Pathways with an FDR lower or equal to 0.05 are represented by -log10 enrichment FDR. **D.** Gene Ontology (GO) enrichment analysis of Cellular Compartment (CC) localization of significant identified proteins upon transient expression of NbSOBIR1-YFP-TbID. All possible localizations for each protein are considered. See Table 1, *Cellular Localization*.

**Table 1.**
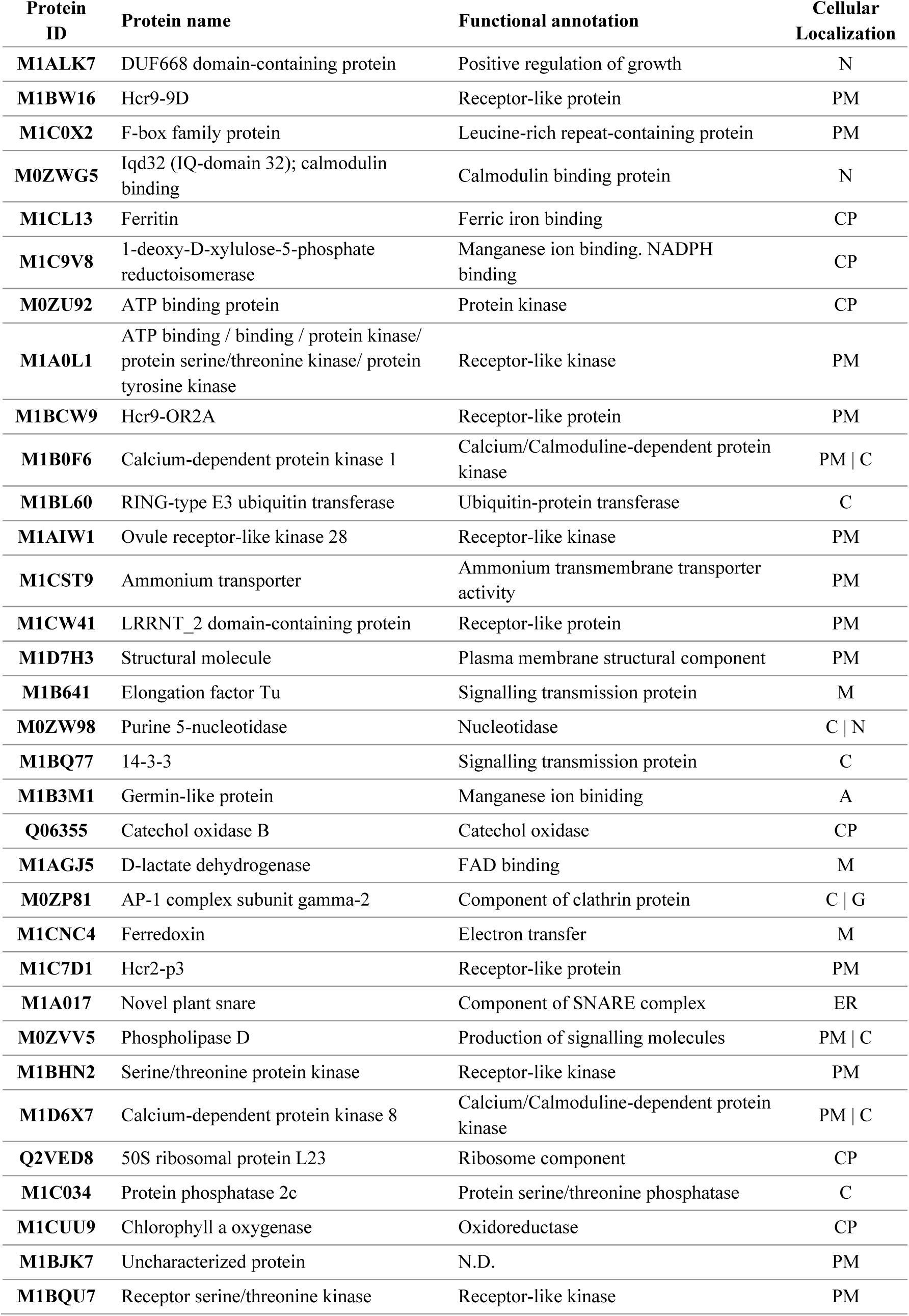
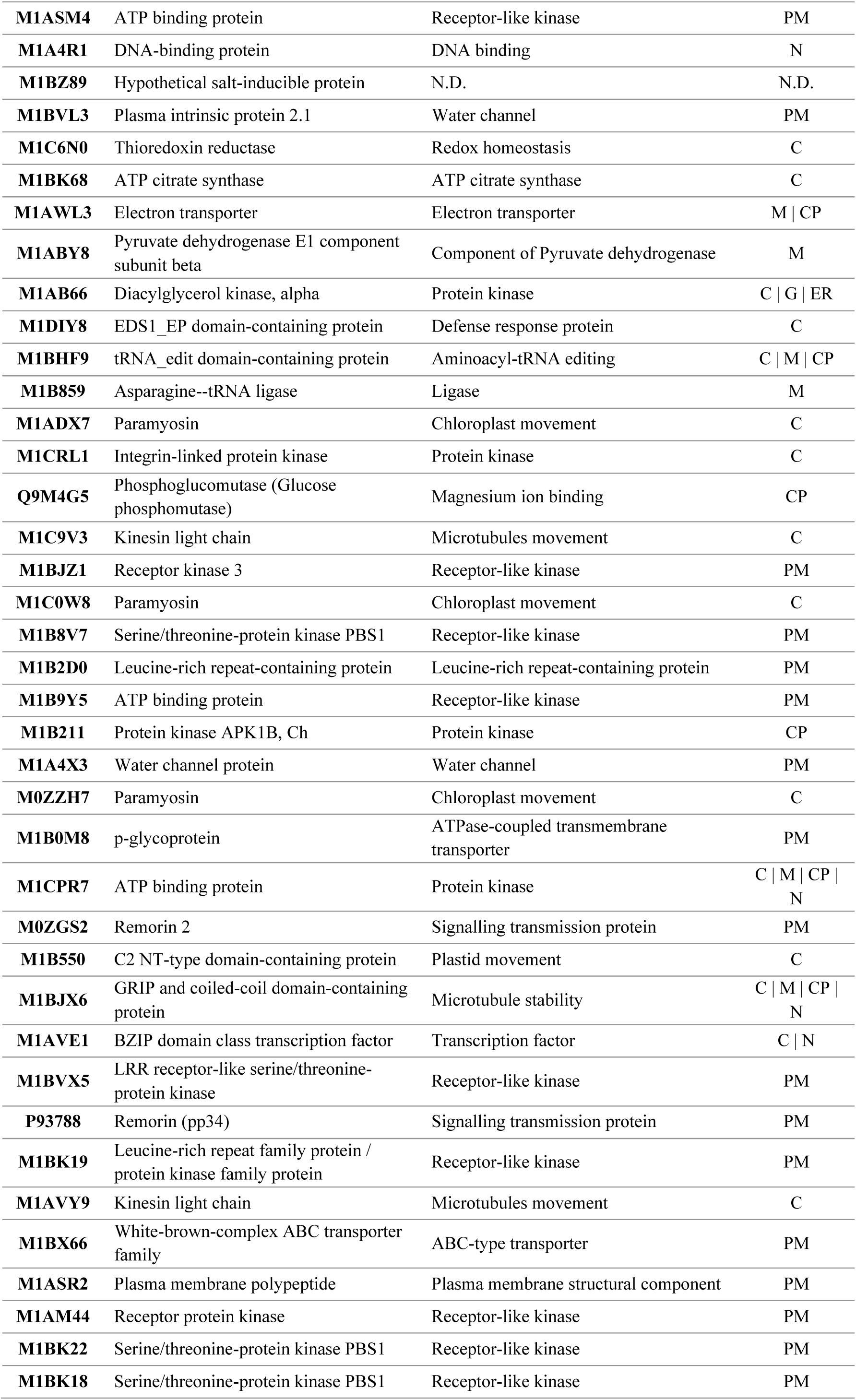
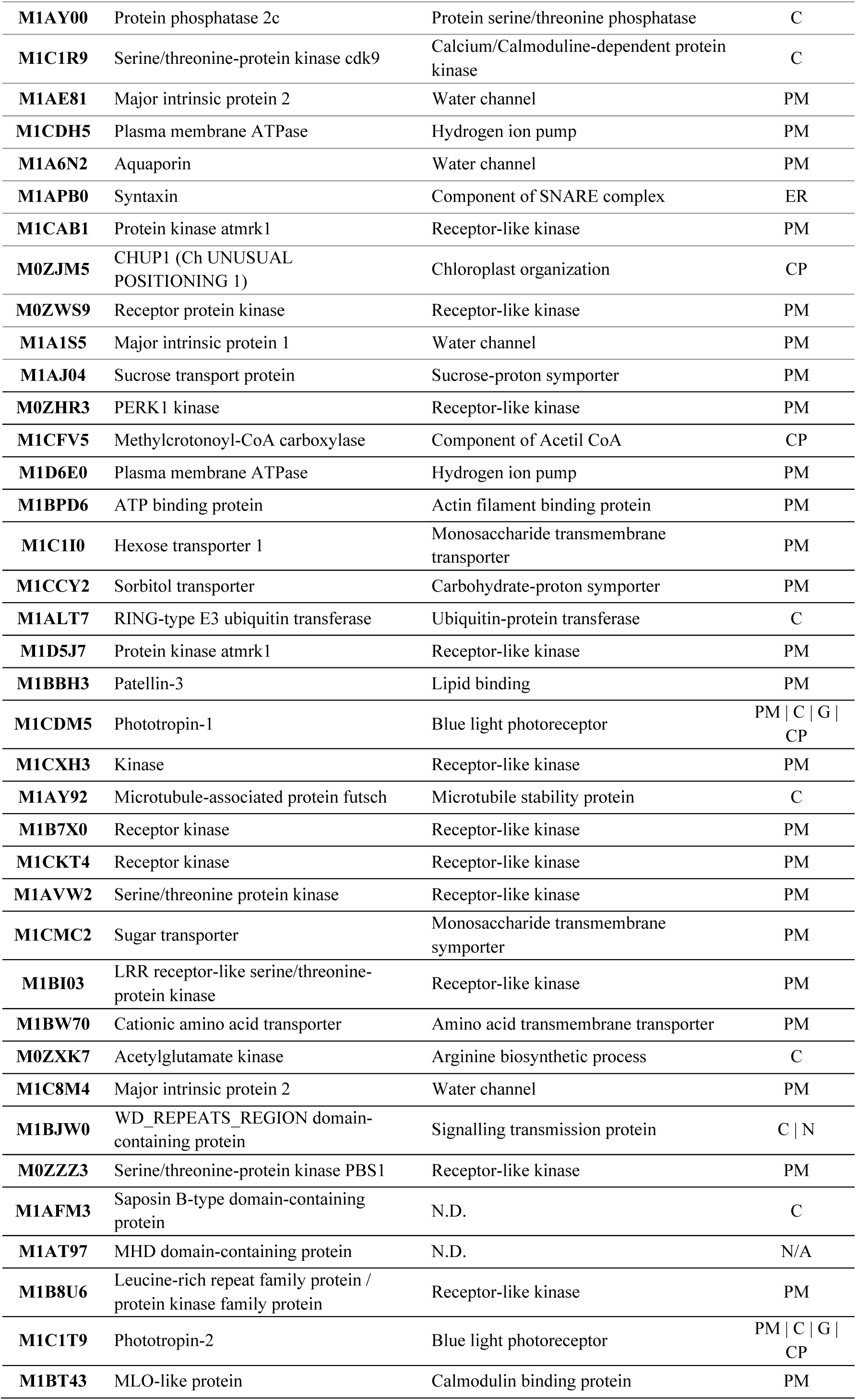
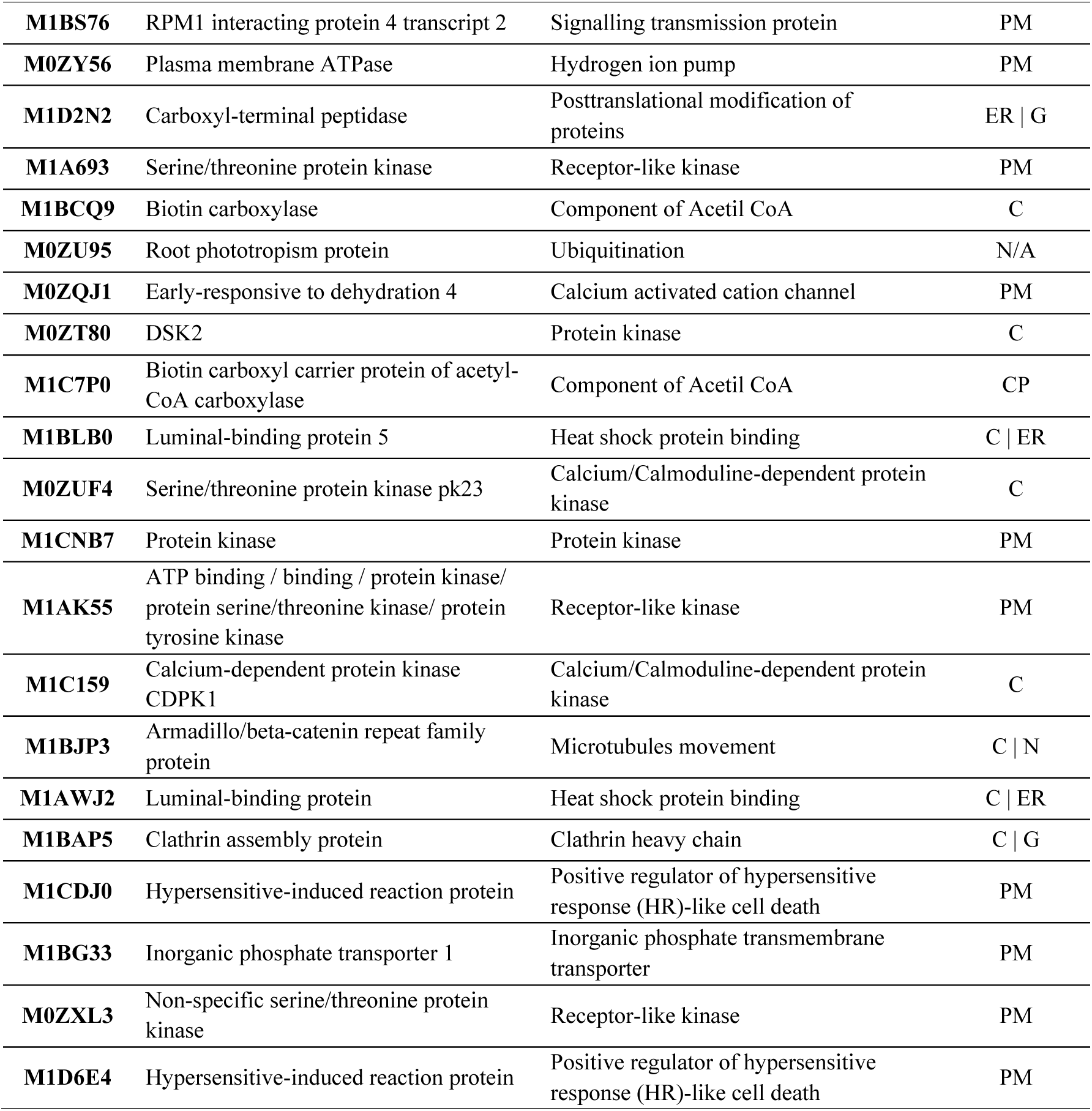
Significant NbSOBIR1 proximal proteins identified by TurboID-MS in NbSOBIR1-YFP-TbID vs AtLTI6B TbID. Protein ID, protein name, functional annotation, and cellular localization were obtained from the UniProt database (UP000011115). When information was not available in UniProt, functional annotation and cellular localization were inferred from homology. Cellular localization: PM, Plasma Membrane; C, Cytosol; CP, Chloroplast; M, Mitochondria; ER, Endoplasmic Reticulum; G: Golgi N, Nucleus; A: Apoplast; N.D.: no data, unknown data or cannot be inferred from homology. Proteins are ranked by FDR enrichment, highest to lowest.

### Elicitation with ELR/INF1 induces a shift in the NbSOBIR1 proxitome

Previous molecular studies demonstrated that ELR constitutively associates with potato SOBIR1 [51]. To investigate dynamic changes in the interactome upon immune activation, we co-expressed NbSOBIR1-YFP-TbID with the receptor ELR and the elicitin INF1 (+E/I) (**Figure 3A**). Comparison of the elicited sample (NbSOBIR1 +E/I) with the AtLTI6B control revealed a markedly altered profile of the NbSOBIR1 proxitome, with only 83 significantly enriched proteins (**Table 2**, **Figure 1C**). Strikingly, ELR and INF1 themselves were not identified among the identified proteins by MS. Furthermore, the functional classification of the interactome shifted dramatically from immune signaling to stress-associated metabolism (**Figure 3B-C**). Enriched proteins included components of the Acetyl-CoA carboxylase complex, chaperones involved in protein folding, and ribosomal proteins. While the majority of candidates were linked to metabolism, several notable exceptions related to pathogen recognition and signal transduction were also captured: the helper NLR, NRC1, which is essential for signaling in Solanaceae [52, 53]; the paralogue RLPs Hcr9ORC2, Hcr9-OR2A, Hcr9-9D and M18S-3Ap [50, 54]; a protein phosphatase 2c (PP2C) and a protein kinase referred to as *ARABIDOPSIS THALIANA* MLK/RAF-RELATED PROTEIN KINASE 1 (ATMRK1), which is a RAF-like kinase homologue from *A. thaliana* involved in various MAPK cascades [55, 56]. Subcellular localization analysis indicated an enrichment of putative interactors localized in the PM (41%) and in the cytoplasm (28%), aligning with previous results (**Figure 3D**).

**Figure 3.**
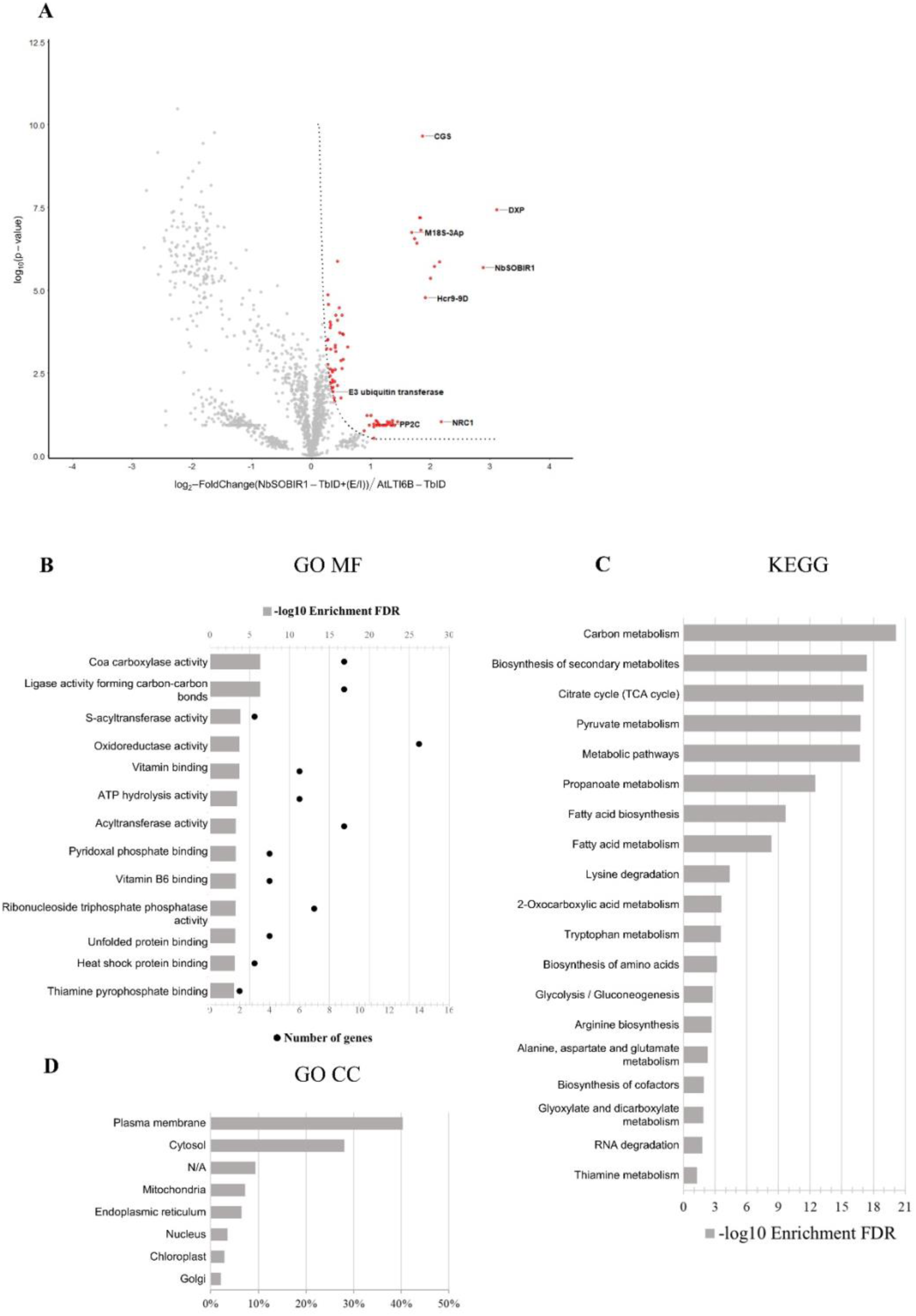
Detecting proteins proximal to NbSOBIR1 in potato MCD321 upon co-infiltration of ELR and INF1. **A.** The Volcano plot depicts the log2-fold change versus the -log2 p-value, with SOBIR1-TbID co-infiltrated with ELR and INF1 (E/I) on the right and AtLTI6B-YFP-TbID on the left. The fold change is the LFQ value of the identified proteins between SOBIR1-TbID with (E/I) and AtLTI6B-YFP-TbID samples. The p-value is the probability of a protein to be significantly proximal to SOBIR1-TbID with (E/I) or to AtLTI6B-YFP-TbID. Significant proteins biotinylated by NbSOBIR1-YFP-TbID were selected using an S0 of 0.1 and a p-value lower or equal to 0.05 (marked by red dots). The grey dots represent non-significant identified proteins with an S0 lower than 0.1 and a p-value higher than 0.05. **B.** Gene Ontology (GO) enrichment analysis of Molecular Function (MF) from significant identified proteins upon transient expression of NbSOBIR1-YFP-TbID co-infiltrated with ELR and INF1. Pathways with an FDR lower or equal to 0.05 are represented by -log10 enrichment FDR as a grey bar. The numbers of genes involved in each pathway are represented by black dots. **C.** KEGG pathway enrichment analysis of significant identified proteins upon transient expression of NbSOBIR1-YFP-TbID co-infiltrated with ELR and INF1. Pathways with an FDR lower or equal to 0,05 are represented by -log10 enrichment FDR. **D.** Gene Ontology (GO) enrichment analysis of Cellular Compartment (CC) localization of significant identified proteins upon transient expression of NbSOBIR1-YFP-TbID co-infiltrated with ELR and INF1. All possible localizations for each protein are considered. See Table 2, *Cellular Localization*.

**Table 2.**
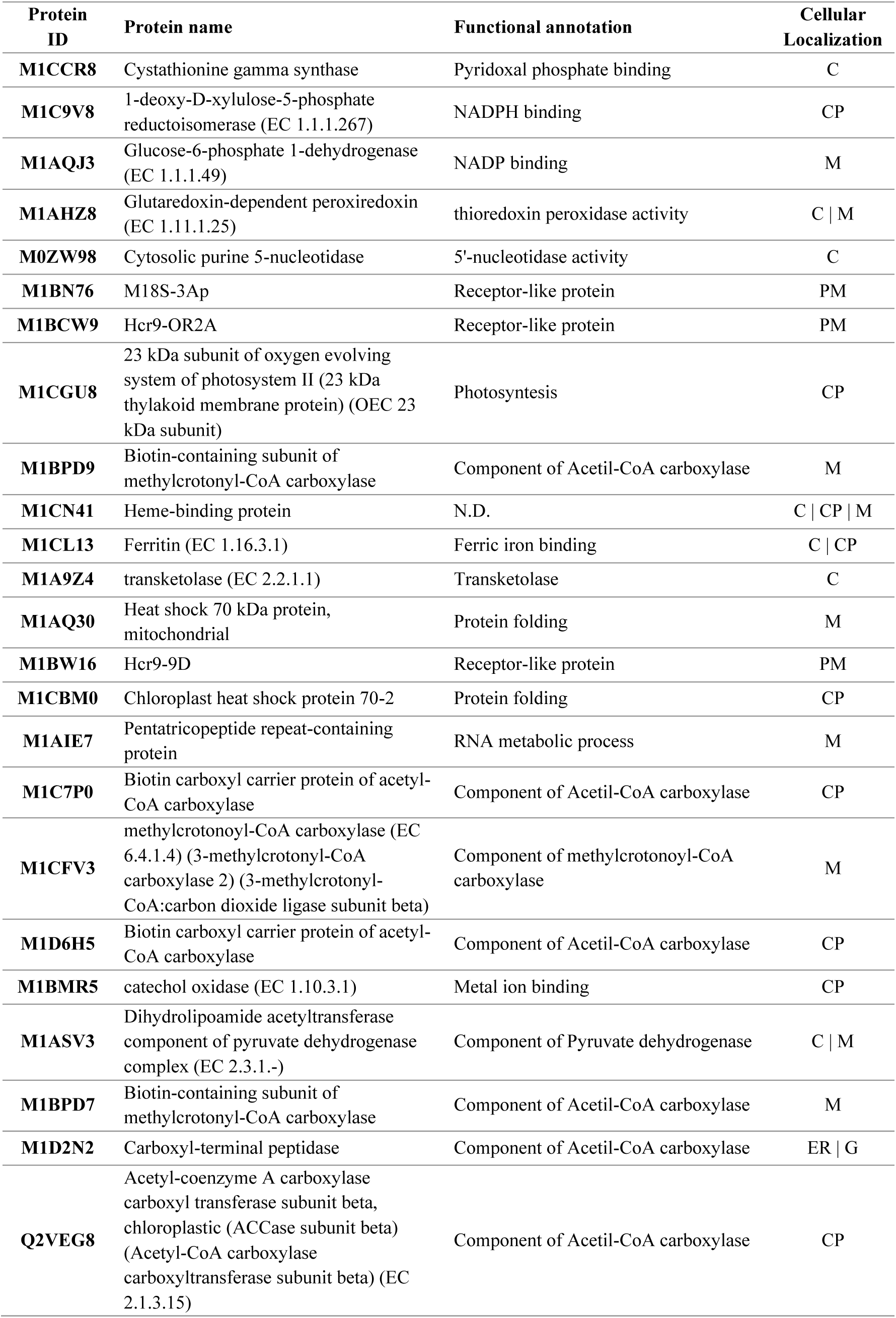
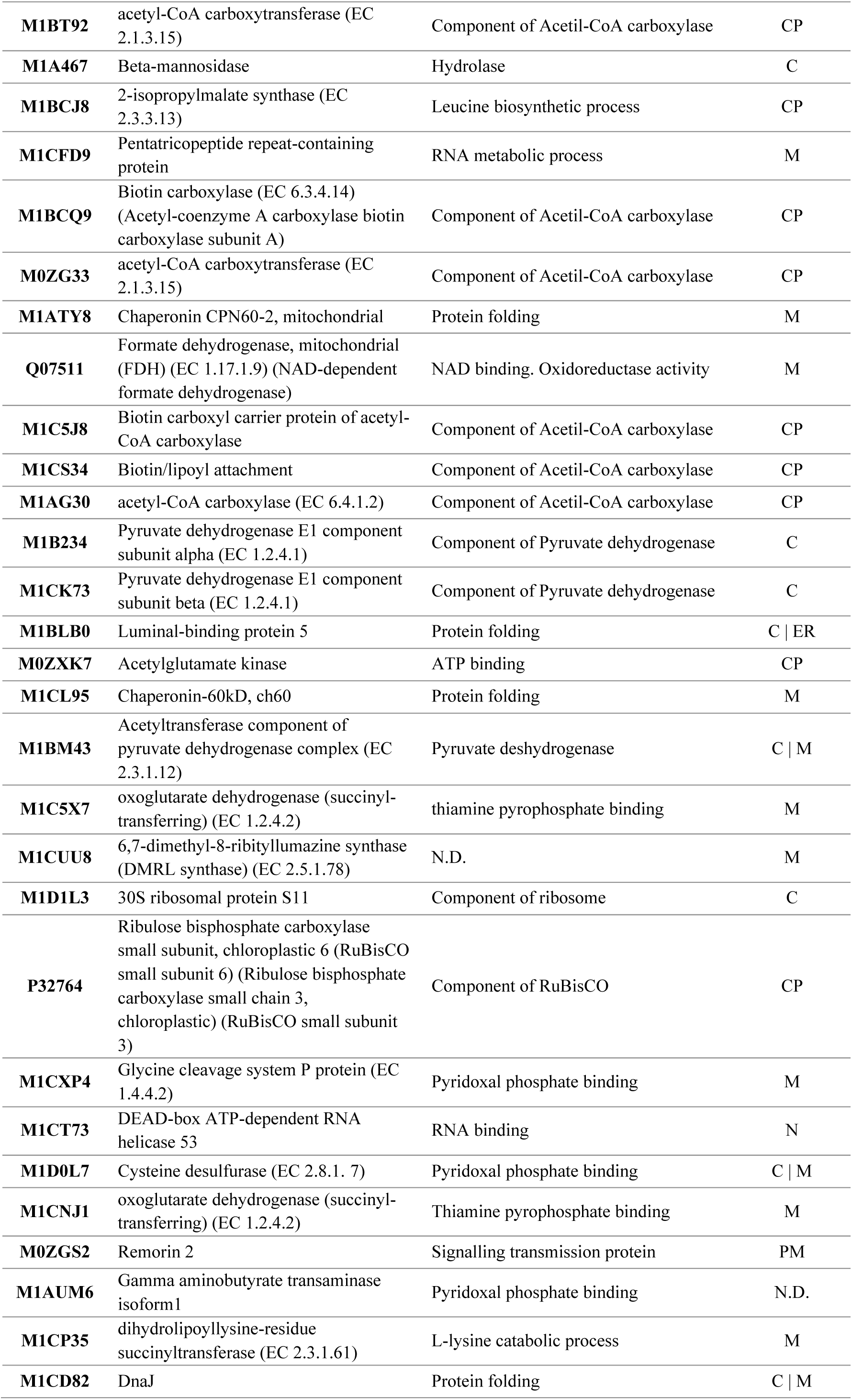
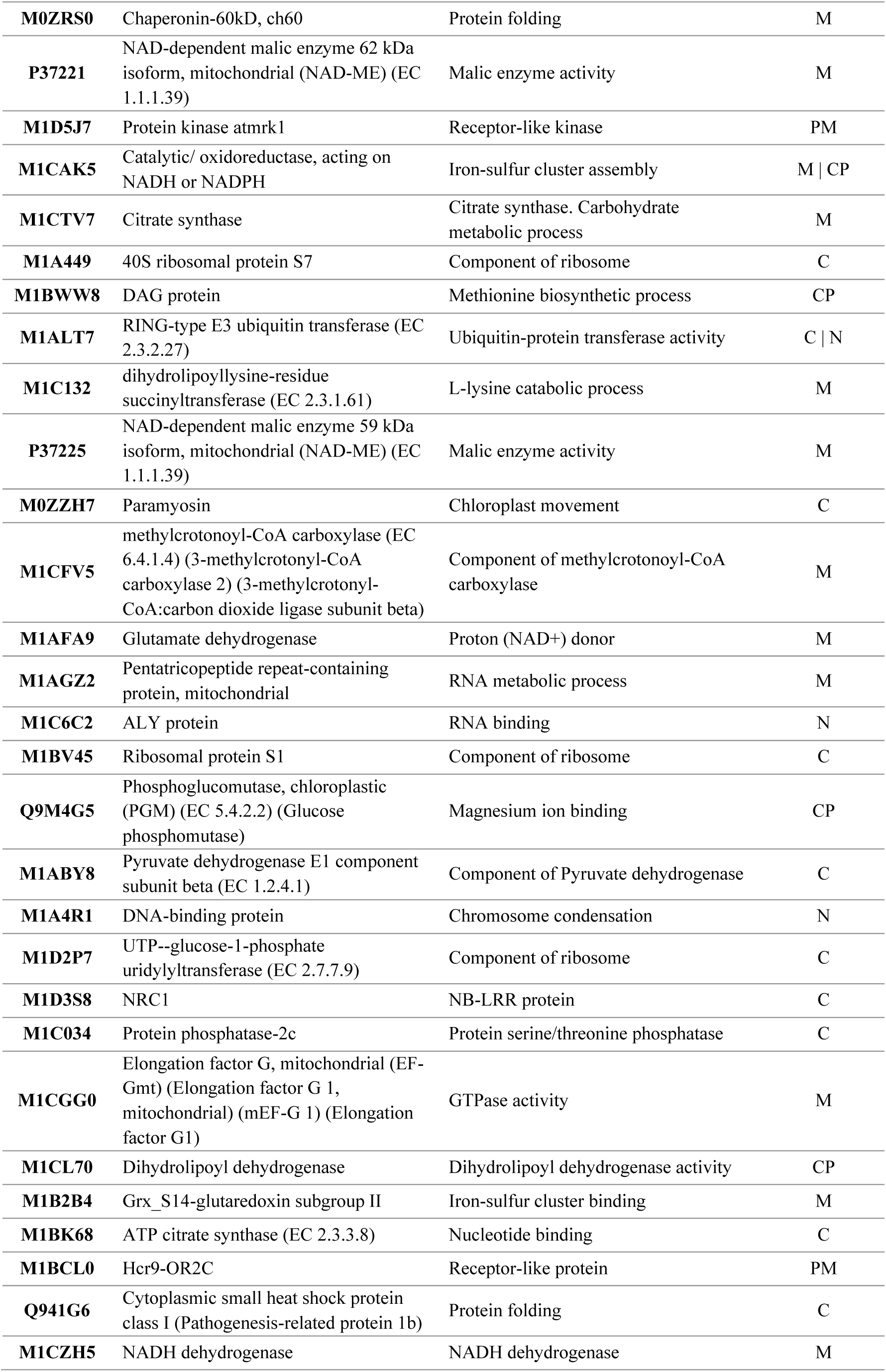
Significant NbSOBIR1 proximal proteins identified upon elicitation by TurboID-MS in NbSOBIR1-YFP-TbID +(E/I) vs AtLTI6B TbID. Protein ID, protein name, functional annotation, and cellular localization were obtained from the UniProt database (UP000011115). When information was not available in UniProt, functional annotation and cellular localization were inferred from homology. Cellular localization: PM, Plasma Membrane; C, Cytosol; CP, Chloroplast; M, Mitochondria; ER, Endoplasmic Reticulum; G: Golgi N, Nucleus; N.D.: no data, unknown data or cannot be inferred from homology. Proteins are ranked by FDR enrichment, highest to lowest.

| **Protein ID** | **Protein name** | **Functional annotation** | **Cellular Localization** |
| --- | --- | --- | --- |
| **M1CCR8** | Cystathionine gamma synthase | Pyridoxal phosphate binding | C |
| **M1C9V8** | 1-deoxy-D-xylulose-5-phosphate reductoisomerase (EC 1.1.1.267) | NADPH binding | CP |
| **M1AQJ3** | Glucose-6-phosphate 1-dehydrogenase (EC 1.1.1.49) | NADP binding | M |
| **M1AHZ8** | Glutaredoxin-dependent peroxiredoxin (EC 1.11.1.25) | thioredoxin peroxidase activity | C \| M |
| **M0ZW98** | Cytosolic purine 5-nucleotidase | 5'-nucleotidase activity | C |
| **M1BN76** | M18S-3Ap | Receptor-like protein | PM |
| **M1BCW9** | Hcr9-OR2A | Receptor-like protein | PM |
| **M1CGU8** | 23 kDa subunit of oxygen evolving system of photosystem II (23 kDa thylakoid membrane protein) (OEC 23 kDa subunit) | Photosynthesis | CP |
| **M1BPD9** | Biotin-containing subunit of methylcrotonyl-CoA carboxylase | Component of Acetil-CoA carboxylase | M |
| **M1CN41** | Heme-binding protein | N.D. | C \| CP \| M |
| **M1CL13** | Ferritin (EC 1.16.3.1) | Ferric iron binding | C \| CP |
| **M1A9Z4** | transketolase (EC 2.2.1.1) | Transketolase | C |
| **M1AQ30** | Heat shock 70 kDa protein, mitochondrial | Protein folding | M |
| **M1BW16** | Hcr9-9D | Receptor-like protein | PM |
| **M1CBM0** | Chloroplast heat shock protein 70-2 | Protein folding | CP |
| **M1AIE7** | Pentatricopeptide repeat-containing protein | RNA metabolic process | M |
| **M1C7P0** | Biotin carboxyl carrier protein of acetyl-CoA carboxylase | Component of Acetil-CoA carboxylase | CP |
| **M1CFV3** | methylcrotonoyl-CoA carboxylase (EC 6.4.1.4) (3-methylcrotonyl-CoA carboxylase 2) (3-methylcrotonyl-CoA:carbon dioxide ligase subunit beta) | Component of methylcrotonoyl-CoA carboxylase | M |
| **M1D6H5** | Biotin carboxyl carrier protein of acetyl-CoA carboxylase | Component of Acetil-CoA carboxylase | CP |
| **M1BMR5** | catechol oxidase (EC 1.10.3.1) | Metal ion binding | CP |
| **M1ASV3** | Dihydrolipoamide acetyltransferase component of pyruvate dehydrogenase complex (EC 2.3.1.-) | Component of Pyruvate dehydrogenase | C \| M |
| **M1BPD7** | Biotin-containing subunit of methylcrotonyl-CoA carboxylase | Component of Acetil-CoA carboxylase | M |
| **M1D2N2** | Carboxyl-terminal peptidase | Component of Acetil-CoA carboxylase | ER \| G |
| **Q2VEG8** | Acetyl-coenzyme A carboxylase carboxyl transferase subunit beta, chloroplastic (ACCase subunit beta) (Acetyl-CoA carboxylase carboxyltransferase subunit beta) (EC 2.1.3.15) | Component of Acetil-CoA carboxylase | CP |
| **M1BT92** | acetyl-CoA carboxytransferase (EC 2.1.3.15) | Component of Acetyl-CoA carboxylase | CP |
| **M1A467** | Beta-mannosidase | Hydrolase | C |
| **M1BCJ8** | 2-isopropylmalate synthase (EC 2.3.3.13) | Leucine biosynthetic process | CP |
| **M1CFD9** | Pentatricopeptide repeat-containing protein | RNA metabolic process | M |
| **M1BCQ9** | Biotin carboxylase (EC 6.3.4.14) (Acetyl-coenzyme A carboxylase biotin carboxylase subunit A) | Component of Acetyl-CoA carboxylase | CP |
| **M0ZG33** | acetyl-CoA carboxytransferase (EC 2.1.3.15) | Component of Acetyl-CoA carboxylase | CP |
| **M1ATY8** | Chaperonin CPN60-2, mitochondrial | Protein folding | M |
| **Q07511** | Formate dehydrogenase, mitochondrial (FDH) (EC 1.17.1.9) (NAD-dependent formate dehydrogenase) | NAD binding. Oxidoreductase activity | M |
| **M1C5J8** | Biotin carboxyl carrier protein of acetyl-CoA carboxylase | Component of Acetyl-CoA carboxylase | CP |
| **M1CS34** | Biotin/lipoyl attachment | Component of Acetyl-CoA carboxylase | CP |
| **M1AG30** | acetyl-CoA carboxylase (EC 6.4.1.2) | Component of Acetyl-CoA carboxylase | CP |
| **M1B234** | Pyruvate dehydrogenase E1 component subunit alpha (EC 1.2.4.1) | Component of Pyruvate dehydrogenase | C |
| **M1CK73** | Pyruvate dehydrogenase E1 component subunit beta (EC 1.2.4.1) | Component of Pyruvate dehydrogenase | C |
| **M1BLB0** | Luminal-binding protein 5 | Protein folding | C \| ER |
| **M0ZXK7** | Acetylglutamate kinase | ATP binding | CP |
| **M1CL95** | Chaperonin-60kD, ch60 | Protein folding | M |
| **M1BM43** | Acetyltransferase component of pyruvate dehydrogenase complex (EC 2.3.1.12) | Pyruvate deshydrogenase | C \| M |
| **M1C5X7** | oxoglutarate dehydrogenase (succinyl-transferring) (EC 1.2.4.2) | thiamine pyrophosphate binding | M |
| **M1CUU8** | 6,7-dimethyl-8-ribityllumazine synthase (DMRL synthase) (EC 2.5.1.78) | N.D. | M |
| **M1D1L3** | 30S ribosomal protein S11 | Component of ribosome | C |
| **P32764** | Ribulose biphosphate carboxylase small subunit, chloroplastic 6 (RuBisCO small subunit 6) (Ribulose biphosphate carboxylase small chain 3, chloroplastic) (RuBisCO small subunit 3) | Component of RuBisCO | CP |
| **M1CXP4** | Glycine cleavage system P protein (EC 1.4.4.2) | Pyridoxal phosphate binding | M |
| **M1CT73** | DEAD-box ATP-dependent RNA helicase 53 | RNA binding | N |
| **M1D0L7** | Cysteine desulfurase (EC 2.8.1. 7) | Pyridoxal phosphate binding | C \| M |
| **M1CNJ1** | oxoglutarate dehydrogenase (succinyl-transferring) (EC 1.2.4.2) | Thiamine pyrophosphate binding | M |
| **M0ZGS2** | Remorin 2 | Signalling transmission protein | PM |
| **M1AUM6** | Gamma aminobutyrate transaminase isoform1 | Pyridoxal phosphate binding | N.D. |
| **M1CP35** | dihydrolipoyllysine-residue succinyltransferase (EC 2.3.1.61) | L-lysine catabolic process | M |
| **M1CD82** | DnaJ | Protein folding | C \| M |
| **M0ZRS0** | Chaperonin-60kD, ch60 | Protein folding | M |
| **P37221** | NAD-dependent malic enzyme 62 kDa isoform, mitochondrial (NAD-ME) (EC 1.1.1.39) | Malic enzyme activity | M |
| **M1D5J7** | Protein kinase atmrk1 | Receptor-like kinase | PM |
| **M1CAK5** | Catalytic/ oxidoreductase, acting on NADH or NADPH | Iron-sulfur cluster assembly | M \| CP |
| **M1CTV7** | Citrate synthase | Citrate synthase. Carbohydrate metabolic process | M |
| **M1A449** | 40S ribosomal protein S7 | Component of ribosome | C |
| **M1BWW8** | DAG protein | Methionine biosynthetic process | CP |
| **M1ALT7** | RING-type E3 ubiquitin transferase (EC 2.3.2.27) | Ubiquitin-protein transferase activity | C \| N |
| **M1C132** | dihydrolipoyllysine-residue succinyltransferase (EC 2.3.1.61) | L-lysine catabolic process | M |
| **P37225** | NAD-dependent malic enzyme 59 kDa isoform, mitochondrial (NAD-ME) (EC 1.1.1.39) | Malic enzyme activity | M |
| **M0ZZH7** | Paramyosin | Chloroplast movement | C |
| **M1CFV5** | methylcrotonoyl-CoA carboxylase (EC 6.4.1.4) (3-methylcrotonoyl-CoA carboxylase 2) (3-methylcrotonyl-CoA:carbon dioxide ligase subunit beta) | Component of methylcrotonoyl-CoA carboxylase | M |
| **M1AFA9** | Glutamate dehydrogenase | Proton (NAD<sup>+</sup>) donor | M |
| **M1AGZ2** | Pentatricopeptide repeat-containing protein, mitochondrial | RNA metabolic process | M |
| **M1C6C2** | ALY protein | RNA binding | N |
| **M1BV45** | Ribosomal protein S1 | Component of ribosome | C |
| **Q9M4G5** | Phosphoglucomutase, chloroplastic (PGM) (EC 5.4.2.2) (Glucose phosphomutase) | Magnesium ion binding | CP |
| **M1ABY8** | Pyruvate dehydrogenase E1 component subunit beta (EC 1.2.4.1) | Component of Pyruvate dehydrogenase | C |
| **M1A4R1** | DNA-binding protein | Chromosome condensation | N |
| **M1D2P7** | UTP--glucose-1-phosphate uridylyltransferase (EC 2.7.7.9) | Component of ribosome | C |
| **M1D3S8** | NRC1 | NB-LRR protein | C |
| **M1C034** | Protein phosphatase-2c | Protein serine/threonine phosphatase | C |
| **M1CGG0** | Elongation factor G, mitochondrial (EF-Gmt) (Elongation factor G 1, mitochondrial) (mEF-G 1) (Elongation factor G1) | GTPase activity | M |
| **M1CL70** | Dihydrolipoyl dehydrogenase | Dihydrolipoyl dehydrogenase activity | CP |
| **M1B2B4** | Grx_S14-glutaredoxin subgroup II | Iron-sulfur cluster binding | M |
| **M1BK68** | ATP citrate synthase (EC 2.3.3.8) | Nucleotide binding | C |
| **M1BCL0** | Hcr9-OR2C | Receptor-like protein | PM |
| **Q941G6** | Cytoplasmic small heat shock protein class I (Pathogenesis-related protein 1b) | Protein folding | C |
| **M1CZH5** | NADH dehydrogenase | NADH dehydrogenase | M |

Since the wild potato genotype MCD-321 lacks the ELR receptor, we hypothesized that the transient co-expression of ELR with INF1 would specifically recruit a unique set of signaling partners to the NbSOBIR1 complex. To dissect these specific dynamic changes, we performed a direct comparative analysis between the elicited (NbSOBIR1 +E/I) and the non-elicited (NbSOBIR1) samples. This analysis resulted in 12 proteins with a significant fold change in the MS dataset of the elicited sample NbSOBIR1-YFP-TbID +E/I **(Figure 4A**, **Table 3-4**). These proteins are predominantly related to changes in the metabolism, including CoA carboxylase subunits (3), cytochrome (1), nucleosidase (1), succinyl-transferase (3), transketolase (1), acetyltransferase activity (1), protein folding (1), and a pentatricopeptide (1) (**Table 4**). Consistent with this role in metabolism, the majority of these proteins are localized to the mitochondria (M) (62%), followed by the chloroplast (CP) (23%) and cytosol (C) (15%). Notably, in this direct comparison, we failed to identify proteins related to plant-pathogen interactions, or signaling in the dataset originating from the elicited sample (NbSOBIR1 +E/I) dataset (**Figure 4**). Instead, the non-elicited NbSOBIR1-YFP-TbID sample yielded a significantly larger proxitome (140 proteins), with 39.3% of these candidates also found in the NbSOBIR1-YFP-TbID set when compared to AtLTI6B-YFP-TbID (**Table 3**, **Figure 1C**). KEGG pathway analysis indicated that plant-pathogen interaction candidates remained at the highest enriched category in the non-elicited sample (**Figure 4C**). Specific downstream signaling components were enriched exclusively in the non-elicited state, when compared to the elicited sample. We identified an AVR RESISTANCE TO PSEUDOMONAS SYRINGAE PATHOVAR TOMATO (AvrRpt)-cleavage domain-containing protein and PTO-INTERACTING PROTEIN 1 (Pti1), both functionally related to RIN4. In addition, a mitogen-activated protein kinase kinase (MAPKK) and MAP KINASE KINASE KINASE 7 (MAPKKK7) were significantly enriched (**Table 3**). In this direct comparison, the identified proteins of the non-elicited NbSOBIR1-YFP-TbID were primarily localized to the mitochondria (36%), chloroplast (23%), and cytosol (26%), with only a small fraction at the PM (8%) (**Figure 4D**). Thus, core complex components of the NbSOBIR1-containing complex, which are mainly located at the PM and in the cytoplasm, such as RLPs, RLKs and RIN4, appear as common proximal interactors of NbSOBIR1 in the elicited and non-elicited samples.

**Figure 4.**
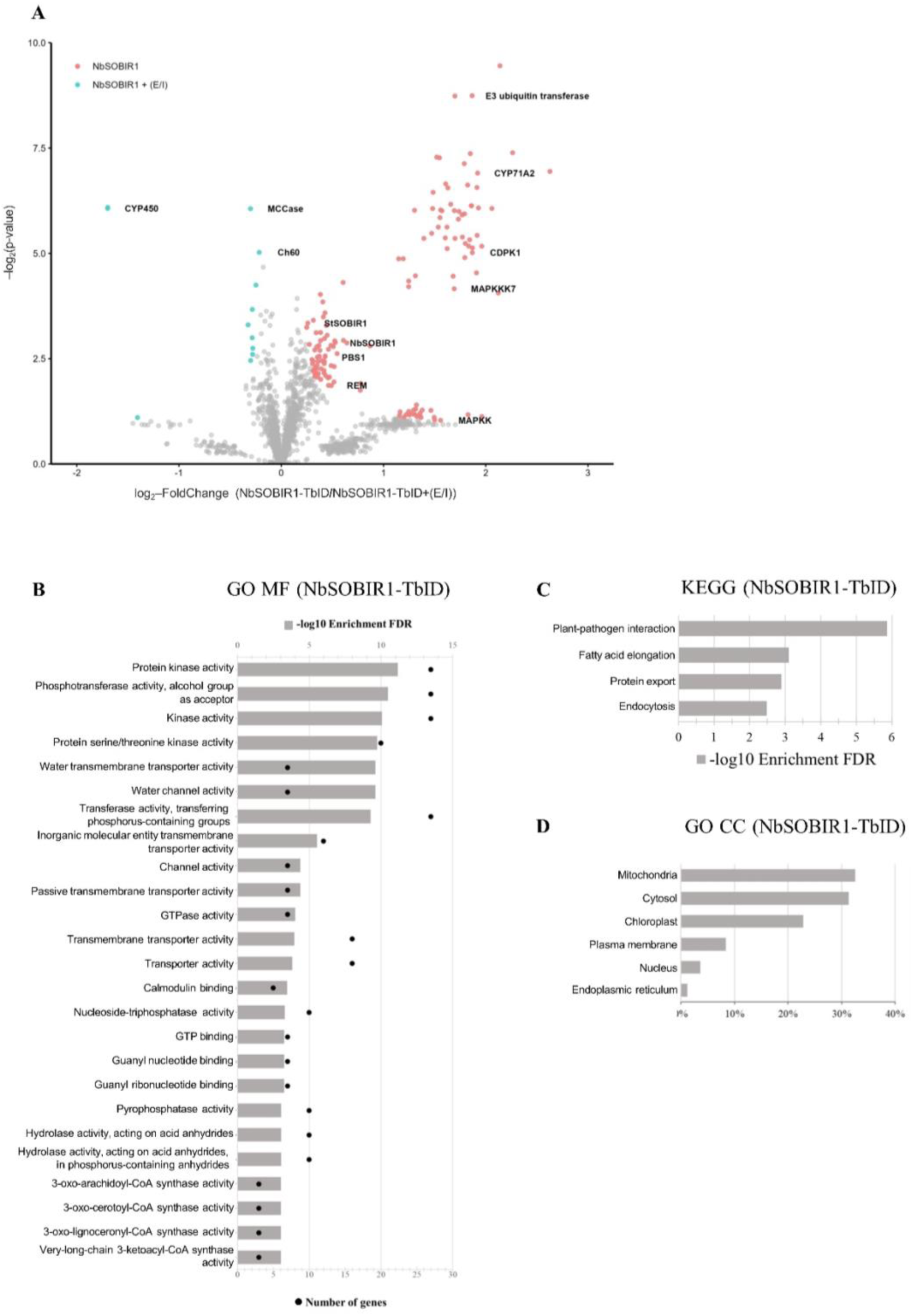
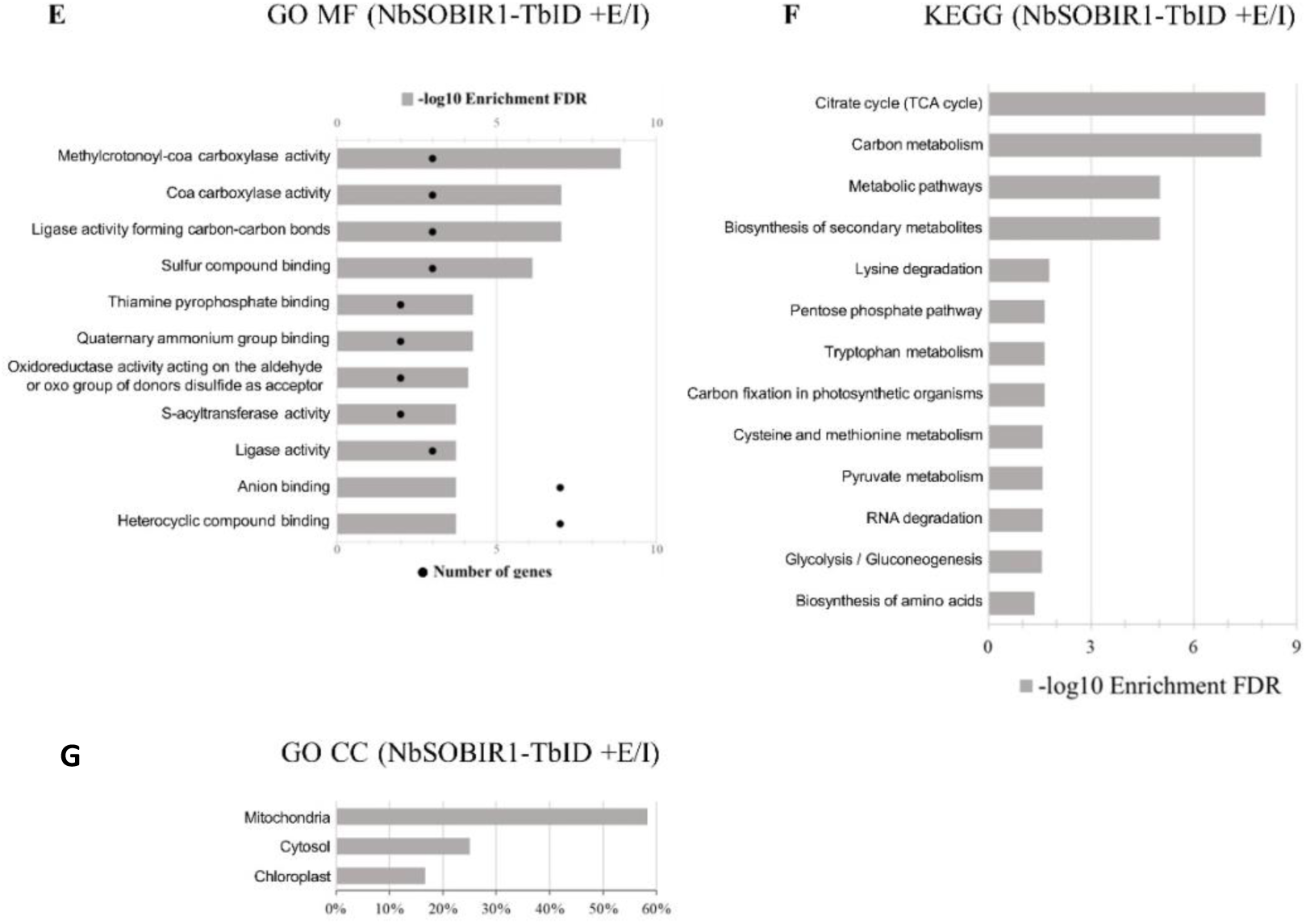
Detecting proteins proximal to NbSOBIR1 in potato MCD321 in presence or absence of ELR and INF1. **A.** The Volcano plot depicts the log2-fold change versus the -log2 p-value, with SOBIR1-TbID on the right and SOBIR1-TbID with ELR and INF1 (E/I) on the left. The fold change is the LFQ value of the identified proteins between SOBIR1-TbID and SOBIR1-TbID with (E/I) samples. The p-value is the probability of a protein to be significantly proximal to SOBIR1-TbID or to SOBIR1-TbID with (E/I). Significant proteins biotinylated by NbSOBIR1-YFP-TbID were selected using an S0 of 0.1 and a p-value lower or equal to 0.05. Significant hits in the presence of ELR and INF1 (E/I) are indicted by blue dots. Significant hits in the absence of ELR and INF1 (E/I) are indicted by red dots. The grey dots represent non-significant identified proteins with an S0 lower than 0.1 and a p-value higher than 0.05. **B, E.** Gene Ontology (GO) enrichment analysis of Molecular Function from significant identified proteins upon transient expression of (**B**) NbSOBIR1-YFP-TbID or co-expression (**E)** with ELR and INF1. Pathways with an FDR lower or equal to 0.05 are represented by -log10 enrichment FDR as a grey bar. Numbers of genes from each pathway are represented by black dots. **C, F.** KEGG pathway enrichment analysis of significant identified proteins upon transient expression of (**E**) NbSOBIR1-YFP-TbID or co-expression (**F)** with ELR and INF1. Pathways with an FDR lower or equal to 0.05 are represented by -log10 enrichment FDR. **D, G.** Gene Ontology (GO) enrichment analysis of Cellular Compartment (CC) localization of significant identified proteins upon transient expression of (**D**) NbSOBIR1-YFP-TbID or co-expression (**G)** with ELR and INF1. All possible localizations for each protein are considered. See Table 3-4, *Cellular Localization*.

**Table 3.** Significant NbSOBIR1 proximal proteins identified by TurboID-MS in NbSOBIR1-YFP-TbID vs A NbSOBIR1-YFP-TbID +(E/I). Protein ID, protein name, functional annotation, and cellular localization were obtained from the UniProt database (UP000011115). When information was not available in UniProt, functional annotation and cellular localization were inferred from homology. Cellular localization: PM, Plasma Membrane; C, Cytosol; CP, Chloroplast; M, Mitochondria; ER, Endoplasmic Reticulum; G: Golgi N, Nucleus; N.D.: no data, unknown data or cannot be inferred from homology. Proteins are ranked by FDR enrichment, highest to lowest.

| **Protein ID** | **Protein name** | **Functional annotation** | **Cellular Localization** |
| --- | --- | --- | --- |
| **M1CAP3** | Lipoxygenase (EC 1.13.11.-) | Metal ion binding | N.D. |
| **M1ALK7** | DUF668 domain-containing protein | N.D. | N.D. |
| **M1C0X2** | F-box family protein | N.D. | PM \| C |
| **Q27S52** | 30S ribosomal protein S14, chloroplastic | rRNA binding | CP |
| **M0ZWG5** | Iqd32 (IQ-domain 32); calmodulin binding | Calcium ion binding | PM \| N |
| **M1AI59** | DUF4149 domain-containing protein | N.D. | N.D. |
| **M0ZRF0** | LysM type receptor kinase | ATP binding | PM |
| **M1A0L1** | ATP binding / binding / protein kinase/ protein serine/threonine kinase/ protein tyrosine kinase | ATP binding | PM |
| **M1B866** | Cytochrome P450 71A2 | Heme binding | M |
| **M1ACB9** | Translation machinery associated TMA7 | N.D. | C \| N |
| **M1CBG8** | zinc_ribbon_10 domain-containing protein | N.D. | ER |
| **M0ZWP6** | Tetratricopeptide repeat protein, tpr | Calcium ion binding | C |
| **D5M912** | Zeaxanthin epoxidase, chloroplastic (EC 1.14.15.21) | FAD binding | CP |
| **M1B0F6** | Calcium-dependent protein kinase 1 | ATP binding | C |
| **M1BL60** | RING-type E3 ubiquitin transferase (EC 2.3.2.27) | Ubiquitin-protein transferase activity | C \| N |
| **M1B865** | Cytochrome P450 71A4 | Heme binding | M |
| **M1AIW1** | Ovule receptor-like kinase 28 | ATP binding | PM |
| **M1CST9** | Ammonium transporter | Ammonium transmembrane transporter activity | PM |
| **M1B3V8** | Ribosomal protein L18ae family | N.D. | ER |
| **M1B1Y8** | Protein kinase family protein | ATP binding | PM |
| **M1CAP7** | Lipoxygenase | Metal ion binding | N.D. |
| **M1ASM4** | ATP binding protein | ATP binding | PM |
| **M1CYJ7** | beta-ketoacyl-[acyl-carrier-protein] synthase I (EC 2.3.1.41) | 3-oxoacyl- | M |
| **M0ZU95** | Root phototropism protein | N.D. | N.D. |
| **Q9M5P8** | Alpha-soluble NSF attachment protein (Alpha-SNAP) (N-ethylmaleimide-sensitive factor attachment protein alpha) | Soluble NSF attachment protein activity | PM \| C |
| **M1B641** | Elongation factor Tu | GTP binding | M |
| **M1CHJ9** | ER membrane protein complex subunit 4 | N.D. | ER |
| **M1DRI9** | Receptor-like serine/threonine-protein kinase (EC 2.7.11.1) | ATP binding | PM |
| **M1CQ41** | Hin1 | N.D. | PM |
| **M1A7H4** | PB1 domain-containing protein | N.D. | C |
| **M1CSE0** | Signal peptidase complex subunit 3 (Microsomal signal peptidase 22 kDa subunit) | N.D. | ER |
| **Q38M52** | Proteasome subunit alpha type | N.D. | C |
| **M1B5K7** | Beta-adaptin-like protein | Clathrin binding | C |
| **M1ACK9** | MAPKKK7 | N.D. | C |
| **M0ZU79** | 4-coumarate:CoA ligase | 4-coumarate-CoA ligase activity | M |
| **M1B8R8** | Ras-related protein RABA1f | GTP binding | C |
| **M1DBH4** | Ran GTPase-activating protein 1 | GTPase activator activity | N |
| **M1CLY9** | Acyl-CoA binding protein | N.D. | M |
| **M0ZVV5** | phospholipase D (EC 3.1.4.4) | Metal ion binding | PM |
| **M1ADI9** | Eukaryotic translation initiation factor 4F / eIF-4F, putative | mRNA binding | C \| N |
| **M1BXT8** | Calcium-transporting ATPase (EC 7.2.2.10) | ATP binding | PM |
| **M1BVQ8** | Bem46 | Palmitoyl-(protein) hydrolase activity | PM \| ER |
| **M0ZIW6** | Signal recognition particle 54 kDa protein | 7S RNA binding | C |
| **M0ZH84** | MAPKK | ATP binding | C |
| **M1A3E5** | RAB11G | GTP binding | C |
| **M1AHC3** | Sorting nexin 3 | Phosphatidylinositol binding | C |
| **M1CGV7** | 3-ketoacyl-CoA synthase (EC 2.3.1.-) | Very-long-chain 3-ketoacyl-CoA synthase activity | M |
| **M1AAU2** | Cytochrome P450 | Heme binding | M |
| **M1BDG2** | AvrRpt-cleavage domain-containing protein | N.D. | PM \| C |
| **M1BLH0** | Ubiquitin carrier protein | ATP binding | N |
| **M0ZII9** | Transmembrane protein 120 homolog | N.D. | PM |
| **M1C883** | Ubiquitin interaction motif-containing protein | N.D. | C \| N |
| **M1B3A3** | Endoplasmic reticulum transmembrane protein | N.D. | ER |
| **M1C5V9** | Ankyrin repeat-containing protein | N.D. | PM |
| **M0ZLX3** | Thioredoxin H | Oxidoreductase activity, acting on a sulfur group of donors, disulfide as acceptor | C |
| **M1BJK7** | Uncharacterized protein | N.D. | PM |
| **M1AQN7** | Secretory carrier-associated membrane protein (Secretory carrier membrane protein) | N.D. | PM |
| **M1AYU6** | Cytohesin 1, 2, 3 | Guanyl-nucleotide exchange factor activity | C |
| **M1AL41** | Calcium ion binding protein | N.D. | N.D. |
| **M1AUT0** | ATP binding protein | ATP binding | PM |
| **M1A6P9** | Early-responsive to dehydration 7 | N.D. | PM |
| **M1BIF4** | Ras GTPase; Sigma-54 factor, interaction region | GTP binding | N.D. |
| **M1C8Q6** | DNA binding protein | Hsp70 protein binding | C |
| **M0ZM12** | SPX, N-terminal; EXS, C-terminal | Inositol hexakisphosphate binding | PM |
| **M1AKC4** | DnaJ protein | N.D. | C \| CP \| N |
| **M0ZU86** | Nucleic acid binding protein | RNA binding | N |
| **M0ZMY5** | 39 kDa EF-Hand containing protein | Calcium ion binding | N.D. |
| **M1AXM2** | Protein transport protein sec16 | N.D. | G |
| **M1CG22** | Protein FAM114A2 | N.D. | N.D. |
| **M1CBT5** | Coiled-coil domain-containing protein 47 | Calcium ion binding | PM |
| **M1AUT7** | DWARF1/DIMINUTO | Delta24-sterol reductase activity | C |
| **M1CG68** | Golgi SNAP receptor complex member 1 | SNAP receptor activity | G |
| **M1B9N6** | Oligosaccharyltransferase complex/magnesium transporter family protein | N.D. | PM |
| **M1B2U5** | NADPH--cytochrome P450 reductase (CPR) (P450R) (EC 1.6.2.4) | Flavin adenine dinucleotide binding | ER |
| **M0ZWH2** | Cytokinin riboside 5'-monophosphate phosphoribohydrolase (EC 3.2.2.n1) | N6-(Delta2-isopentenyl)-adenosine 5'-monophosphate phosphoribohydrolase activity | C |
| **M0ZGX0** | Angustifolia | Kinase binding | C |
| **M1AYV8** | Nucleotide binding protein | N.D. | N.D. |
| **M1BJZ1** | Receptor kinase 3 | ATP binding | PM |
| **M1ADI7** | Signal recognition particle receptor alpha subunit | GTP binding | ER |
| **M1ADX7** | Paramyosin | N.D. | C |
| **M1AHN7** | Arf gtpase-activating protein | GTPase activator activity | C \| G |
| **M1B211** | Protein kinase APK1B, chloroplast | ATP binding | CP |
| **M1BV08** | Aquaporin | Water channel activity | PM |
| **M1B8V7** | Serine/threonine-protein kinase PBS1 | ATP binding | PM |
| **M1AVE1** | BZIP domain class transcription factor | N.D. | C |
| **M1BVL3** | Plasma intrinsic protein 2,1 | Water channel activity | PM |
| **M1C9A7** | Anthranilate N-benzoyltransferase protein | Acyltransferase activity | C |
| **M1A8I4** | ASR4 | N.D. | N |
| **M1CPR7** | ATP binding protein | ATP binding | PM |
| **M1A8I6** | Ci21A protein | N.D. | C |
| **M1C1R9** | Serine/threonine-protein kinase cdk9 | ATP binding | C |
| **M1A4X3** | Water channel protein | Water channel activity | PM |
| **P93788** | Remorin (pp34) | Galacturonate binding | PM |
| **M1C0W8** | Paramyosin | N.D. | C |
| **M1AT67** | Co-chaperone protein p23 | Chaperone binding | C |
| **M1AE81** | Major intrinsic protein 2 | Water channel activity | PM |
| **M1CCY2** | Sorbitol transporter | Carbohydrate:proton symporter activity | PM |
| **M1A6N2** | Aquaporin | Water channel activity | PM |
| **M1BK18** | Serine/threonine-protein kinase PBS1 | ATP binding | PM |
| **M1AFM3** | Saposin B-type domain-containing protein | N.D. | C |
| **M1B7X0** | Receptor kinase | ATP binding | PM |
| **M1A1S5** | Major intrinsic protein 1 | Water channel activity | PM |
| **M1A4N1** | DUF639 domain-containing protein | N.D. | N.D. |
| **M1BK19** | Leucine-rich repeat family protein / protein kinase family protein | ATP binding | PM |
| **M1BJX6** | GRIP and coiled-coil domain-containing protein | N.D. | C \| M \| CP \| N |
| **M1C1T9** | Phototropin-2 | Photoreceptor activity | N |
| **M1ANF4** | Calcium lipid binding protein | Lipid binding | PM |
| **M0ZJM5** | CHUP1 (CHLOROPLAST UNUSUAL POSITIONING 1) | N.D. | CP |
| **M1CX27** | Cellulose synthase CslE | Cellulose synthase (UDP-forming) activity | PM |
| **M1D699** | Protein phosphatase 2c | Protein serine/threonine phosphatase activity | C |
| **B7SSK0** | 3-ketoacyl-CoA synthase (EC 2.3.1.-) | Very-long-chain 3-ketoacyl-CoA synthase activity | M |
| **M0ZGS2** | Remorin 2 | N.D. | PM |
| **M1B550** | C2 NT-type domain-containing protein | Plastid movement | C |
| **M1CDM5** | Phototropin-1 | ATP binding | C |
| **M1D6E0** | Plasma membrane ATPase (EC 7.1.2.1) | ATP binding | PM |
| **M1CT77** | Reticulon-like protein | N.D. | ER |
| **M1C8M4** | Major intrinsic protein 2 | Water channel activity | PM |
| **M1CBR3** | Ras-related protein RABA2a | GTP binding | C |
| **M0ZGA9** | Ethylene-responsive late embryogenesis | N.D. | N.D. |
| **M1B694** | ATSLY1 | Syntaxin binding | G |
| **M1ABK5** | 3-ketoacyl-CoA synthase (EC 2.3.1.-) | Very-long-chain 3-ketoacyl-CoA synthase activity | M |
| **M1CSV4** | Transmembrane protein | N.D. | ER |
| **M1ASR2** | Plasma membrane polypeptide | N.D. | PM |
| **M1C1I0** | Hexose transporter 1 | Monosaccharide transmembrane transporter activity | PM |
| **M1BX66** | White-brown-complex ABC transporter family | ABC-type transporter activity | PM |
| **M1BT43** | MLO-like protein | Calcium ion binding | PM |
| **M1B8U6** | Leucine-rich repeat family protein / protein kinase family protein | ATP binding | PM |
| **M1C159** | Calcium-dependent protein kinase CDPK1 | ATP binding | C |
| **M1A0F2** | Pto-interacting protein 1 | ATP binding | PM \| C |
| **M1AT97** | MHD domain-containing protein | N.D. | N.D. |
| **M0ZZZ3** | Serine/threonine-protein kinase PBS1 | ATP binding | PM |
| **M1CAR8** | Acyl-CoA binding protein | Fatty-acyl-CoA binding | C |
| **M1AQS2** | Lipoxygenase (EC 1.13.11.-) | N.D. | N.D. |
| **M1BW70** | Cationic amino acid transporter | N.D. | N.D. |
| **M1CXH3** | Kinase | N.D. | N.D. |
| **M1CDH5** | Plasma membrane ATPase (EC 7.1.2.1) | N.D. | N.D. |
| **M1AM44** | Receptor protein kinase | N.D. | N.D. |
| **M1CKT4** | Receptor kinase | N.D. | N.D. |
| **M1CNB7** | Protein kinase | N.D. | N.D. |

**Table 4.** Significant NbSOBIR1 proximal proteins identified upon elicitation by TurboID-MS in NbSOBIR1-YFP-TbID +(E/I) vs NbSOBIR1-YFP-TbID. Protein ID, protein name, functional annotation, and cellular localization were obtained from the UniProt database (UP000011115). When information was not available in UniProt, functional annotation and cellular localization were inferred from homology. Cellular localization: PM, Plasma Membrane; C, Cytosol; CP, Chloroplast; M, Mitochondria; ER, Endoplasmic Reticulum; G: Golgi N, Nucleus; N.D.: no data, unknown data or cannot be inferred from homology. Proteins are ranked by FDR enrichment, highest to lowest

| **Protein ID** | **Protein name** | **Functional annotation** | **Cellular Localization** |
| --- | --- | --- | --- |
| **M1CL95** | Chaperonin-60kD, ch60 | ATP binding | M |
| **M1BPD7** | Biotin-containing subunit of methylcrotonyl-CoA carboxylase | N.D. | M |
| **M1C5X7** | Oxoglutarate dehydrogenase (succinyl-transferring) (EC 1.2.4.2) | Oxoglutarate dehydrogenase (succinyl-transferring) activity | M |
| **M1CNJ1** | Oxoglutarate dehydrogenase (succinyl-transferring) (EC 1.2.4.2) | Oxoglutarate dehydrogenase (succinyl-transferring) activity | M |
| **M1CFV3** | Methylcrotonoyl-CoA carboxylase (EC 6.4.1.4) (3-methylcrotonyl-CoA carboxylase 2) (3-methylcrotonyl-CoA:carbon dioxide ligase subunit beta) | ATP binding | CP |
| **M1CP35** | Dihydrolipoyllysine-residue succinyltransferase (EC 2.3.1.61) | Dihydrolipoyllysine-residue succinyltransferase activity | M |
| **M1BM43** | Acetyltransferase component of pyruvate dehydrogenase complex (EC 2.3.1.12) | Acetyltransferase activity | C \| M |
| **M1BPD9** | Biotin-containing subunit of methylcrotonyl-CoA carboxylase | ATP binding | M |
| **M1AIE7** | Pentatricopeptide repeat-containing protein | N.D. | CP |
| **M1A9Z4** | Transketolase (EC 2.2.1.1) | Metal ion binding | C |
| **M0ZMY2** | Mta/sah nucleosidase | Methylthioadenosine nucleosidase activity | C |
| **M1AYX7** | Cytochrome P450 | Heme binding | M |

### Low accumulation levels of NbSOBIR1-YFP-TbID affect the identification of putative interactors

When we analyzed the specific enrichment of putative proximal interactors of the elicited sample (NbSOBIR1 +E/I) against the non-elicited sample (NbSOBIR1), we observed a significant reduction in the detection of key immune components. We hypothesized that the reduction in identified interactors upon elicitation might stem from protein instability or low TurboID biotinylating activity. Analysis of the Label-Free Quantification (LFQ) intensities revealed that the intensities of the peptides originating from NbSOBIR1, YFP, and TbID were three-fold lower in the samples co-expressing ELR and INF1 with NbSOBIR1-YFP-TbID, when compared to transient expression of NbSOBIR1-YFP-TbID alone (**Table S1**). Conversely, the AtLTI6B-YFP-TbID control showed YFP and TbID accumulation levels that were three-fold higher than those observed for non-elicited sample, and nine-fold higher than those in elicited sample (**Table S1**). This reduced accumulation of the bait protein in the elicited sample may correlate with the lower number of identified interactors and the low abundance of immune signaling components in the elicited dataset.

## Discussion

Mapping PPIs in crop plants is critical to understanding how defense-related components exactly function, yet deciphering PPIs remains particularly challenging in non-model species. Traditional methods, such as IPs, are often limited by low protein expression levels, only transient and weak interactions taking place, and by the complex biochemistry of membrane-associated defense-related complexes [57]. In this study, we pioneered the use of TurboID-based PL employing the immune-related protein NbSOBIR1 in a non-model plant, the wild potato *S. microdontum*. After screening a set of wild *Solanum* F1 progeny plants with a *S. microdontum* background, we selected the genotype MCD-321, which proved to be highly amenable to *Agrobacterium*-mediated transient transformation. We preferred this genetic background as this is a valuable source for resistance to the oomycete pathogen *P. infestans* [22, 26], and in this background, the signaling pathway to mount the hypersensitive response (HR) upon agro-co-infiltration of ELR and INF1 is conserved [22, 24]. We successfully expressed functional NbSOBIR1-YFP-TurboID (NbSOBIR1-YFP-TbID) fusion protein, confirming its accumulation *in planta* (**Figure 1**) and enabling the identification of a high-confidence set of putative NbSOBIR1 interactors (**Tables 1-2**). These results align with previous plant PL studies [37, 39, 41, 58], establishing the TurboID-PL approach as a powerful tool for potato immune interactomics.

The RLK SOBIR1 is a key regulator of plant immune signaling, which is essential for the function of numerous cell surface-localized RLP immune receptors [7, 8, 14]. The PL-generated data set revealed that NbSOBIR1-proximal proteins are primarily associated with the core plant-pathogen-related immune response, including RLPs, RLKs, and key signaling components like remorin, RIN4, and CDPKs (**Figure 2**). The identification of RLPs related to the *C. fulvum* resistance protein family of tomato (Hcr9, Hcr2 and HcrVf2 paralogues) proves the biological relevance of our dataset, confirming the role of SOBIR1 as a general adaptor of resistance-related RLPs [7, 51, 59, 60]. Our analysis also identified a protein containing an ENHANCED DISEASE SUSCEPTIBILITY 1 (EDS1) domain, presumably representing a functional Solanum ortholog of the EDS1 family (**Table 1**). These findings align with the reported association of EDS1-PHYTOALEXIN-DEFICIENT 4 (PAD4) with SOBIR1 in *Arabidopsis* [61]. We identified four homologues of AVRPPHB SENSITIVE 1 (PBS1), which belong to the class VII RLCKs, which are closely related to BIK1. PBS1 is a known key component of the immune signaling network, and is required for RESISTANCE TO *PSEUDOMONAS SYRINGAE* 5 (RPS5)-mediated HR in *Arabidopsis* [62] and potato [63], but not yet described to interact with SOBIR1. Similarly, NbSOBIR1 was found to be in proximity to class VII RLCKs from *N. benthamiana* and the same study reported that a group of tomato class VII RLCKs physically interact with SlSOBIR1 [14]. This finding extends to other novel kinase candidates in our dataset, like the AtMRK1 homologue, suggesting that NbSOBIR1 is indeed involved in downstream signaling initiated by ligand binding and subsequent BAK1 recruitment by the associated RLP [64]. The identification of a 14-3-3 protein alongside these kinases provides a plausible molecular mechanism for signaling regulation, as these proteins are involved in signal transduction by inducing conformational changes in phosphorylated targets [65–67]. Additionally, components of the SOLUBLE N-ETHYLMALEIMIDE-SENSITIVE FACTOR ATTACHMENT PROTEIN RECEPTOR (SNARE) complex and clathrin were identified in the dataset, suggesting the involvement of vesicular trafficking and endocytosis in receptor recycling and immune regulation [13, 68, 69]. Collectively, our findings suggest the evolutionary conservation of core signaling components that participate in different immune pathways across plant species and reveal a new set of candidates potentially required for the functionality of SOBIR1.

Elicitation by transiently expressing the immune receptor ELR in combination with INF1 elicitin altered the proxitome profile, shifting it towards proteins associated with primary metabolism (e. g. acetyl-CoA carboxylase and ribosomal proteins) and showing a reduction in the number of defense-related putative interactors (**Figure 3**). ELR and INF1 were not detected in the proxitome, and furthermore, we observed a three-fold decrease in the number of NbSOBIR1-derived peptides (**Table S1**). Co-expression of ELR and INF1 triggers an HR (**Figure 1, Figure S2**), which involves an upregulation of cell death-associated proteins and likely contributes to the significantly reduced accumulation of signaling components observed during elicitation (**Figure 3**). Moreover, the continuous induction of HR involves the subsequent endocytosis and degradation of the activated ELR/NbSOBIR1-YFP-TbID complex, which detection could be affected by the rapid turnover of ELR. Studies on the Cf-4/SlSOBIR1 complex have shown that ligand perception triggers rapid co-internalization and degradation of the complex [13]. Similar to the *Arabidopsis* FLS2 receptor, which is targeted to ubiquitin-mediated degradation by E3 ligases upon ligand perception to attenuate immune signaling [70]. The active degradation mechanism might explain the significant reduced accumulation of NbSOBIR1-YFP-TbID in the elicited samples. Consequently, resulting in the loss of lower-abundance or transient putative interactors and the preferential detection of highly abundant enzymes involved in the altered metabolism that may compete for biotin or bind streptavidin non-specifically. To circumvent this receptor turnover, applying the elicitor protein (INF1) directly to the leaf tissue shortly before sample collection, may allow the pull down of transient signaling complexes prior to their endocytic degradation.

The RLK BAK1/SERK3, which is recruited by the RLP/SOBIR1 complex upon its activation, was not identified in our statistical analysis, although it was listed within the non-significantly identified interactors (**Table S2**). The absence of known interactors was also observed in other PL studies [39, 41, 71]. These observations highlight that while informative, the PL approach has its limitations. Still, we captured NRC1, a cytosolic CC-NB-LRR helper NLR. This finding, if confirmed, would provide novel evidence that SOBIR1-dependent surface receptors directly tap into the intracellular NRC network to execute cell death. This observation is in line with the finding that NRC1 and NRC3 act as a node downstream of Cf-4, triggering HR upon recognition of Avr4and thereby connect RLP/SOBIR1-triggered immune initiation with the intracellular immune signaling network in Solanaceae [53, 72].

The specificity of the MS dataset relies on stringent controls to eliminate false positives. While our data validated the PM localization of NbSOBIR1 [7, 8, 13], the AtLTI6B control accumulated TbID levels three times higher than the bait. This imbalance risks non-specific biotinylation, potentially masking specific interactions of NbSOBIR1 with proteins, such as BAK1/SERK3. Similarly, the enrichment of endogenous biotin-containing carboxylases [73] highlights the need for additional controls, such as an untagged bait [58] or using the PM-marker AUTOINHIBITED CA^2+^-ATPASE (ACA8) tagged with TbID, to ensure the stringency in defense network mapping [13, 74]. Optimization of experimental parameters also remains critical [33, 37, 39, 41]; although we utilized limited amounts of additional biotin and proteasome inhibitors, bait accumulation was variable. Future refinements could involve alternative *Agrobacterium* strains for transient expression [75], the use of silencing inhibitors, such as P19 [76] and/or organelle fractionation, such as PM isolation. Finally, data interpretation is hampered by the still incomplete potato reference proteome. Only a few potato varieties have been fully sequenced and annotated [55–57], but these annotations are inferred from model plants, and unique peptides often map ambiguously to multiple proteins. Despite these constraints, the successful detection of previously known SOBIR1 interactors and signaling components validates the robustness of our PL approach.

In conclusion, TurboID-based PL in leaves of potato is a useful technique to decipher the composition of PM-associated receptor complexes. The experimental design is crucial to identify specific interactions with the bait protein of interest, especially with the selection of the right controls to mitigate expression imbalances. The putative interactors identified in this study require further validation through complementary techniques such as gene silencing (VIGS), knock-out (CRISPR/Cas) and/or co-immunoprecipitations. As the capability to engineer immune receptors expands, the integration of identified host targets as sensor domains for effectors has emerged as a powerful strategy for creating new disease resistance specificities. Ultimately, continued investment in open experimental proteome databases for diverse crop species is indispensable to fully leverage the molecular network data generated by PL and to advance crop disease resistance research.

## Materials and methods

### Plant materials and growth conditions

*Solanum* genotypes used for HR screening are listed in **Figure S2**. Progeny genotypes, including MCD-321, were generated from a cross between *S. microdontum* (MCD360-1) and the 3341-15 F1 clone from *S. microdontum* ssp*. gigantophyllum* (GIG362-6) x *S. verrucosum* 3316-17 [24]. MCD360-1 and GIG362-6 were also included in the screening. Plants were maintained and propagated as described before [45]. Two weeks old plantlets were transferred to soil in a climate-controlled greenhouse under long day conditions (16 h-day/8 h-dark), at 18 - 22°C and 70% relative humidity. First, plants were grown for one week in pots with a volume of 0.08 L (5 cm in diameter), then transferred to larger cylindrical pots of 1.7 L (14 cm in diameter) and grown for an additional 2-3 weeks, after which the *Agrobacterium tumefaciens* transient transformation assays were performed.

### Constructs for *Agrobacterium tumefaciens*-mediated transient transformation

The constructs INF1-pK7WG2, ELR-pK7WG2 and empty vector (EV)-pK7WG2 transformed into *A. tumefaciens* strain AGL1 were previously described [21]. NbSOBIR1-TurboID-pEarleyGate101 (NbSOBIR1-YFP-TbID) and AtLTI6B-TurboID-pEarleyGate101 (AtLTI6B-YFP-TbID), transformed into *A. tumefaciens* strain C58C1 were obtained from the Laboratory of Phytopathology (Wageningen University & Research, the Netherlands) [14]. The TbID constructs are C-terminally fused to the TurboID moiety (35 kDa) and the yellow fluorescent protein (YFP) (27 kDa), which results in a full protein of 136 kDa for NbSOBIR1-YFP-TbID, and of 70 kDa for AtLTI6B (**Figure S1**).

### *Agrobacterium tumefaciens*-mediated transient transformation assay

Agroinfiltrations were performed as described [45], using an OD_600_ of 0.3. For the *Solanum* screening, pK7WG2 constructs were agro-infiltrated, or agro-co-infiltrated at 1:1 ratio, in three fully expanded leaves of three plants, in at least two independent experiments. The screening for an HR was evaluated at 5 dpi. For the evaluation of the expression levels of the TbID-fused proteins by western blotting, two fully expanded leaves of two MCD-321 potato plants were agro-infiltrated per construct. For PL experiments, three fully expanded leaves of five MCD-321 potato plants were agro-infiltrated per construct, or agro-co-infiltrated in a 1:1:1 ratio. Each PL sample was prepared in triplicate for eventual LC-MS analysis.

### Total protein extraction and enrichment of biotinylated proteins

Extraction and enrichment of biotinylated proteins was performed as described earlier [57]. Briefly, two days post agro-infiltration, leaves for western blot analysis and PL experiments were infiltrated with a biotin solution (200 µM biotin, 10 mM MES, pH 8), supplemented with 40 µM proteasome inhibitor MG-132 (Sigma Aldrich). Leaf material was harvested at 1 hour post infiltration of the biotin solution and was ground to a fine powder in liquid nitrogen, using a mortar and pestle. The powder was resuspended in ice-cold modified radio-immunoprecipitation assay (RIPA) extraction buffer (Tris-HCl 25 mM, pH 7.6, NaCl 150 mM, IPEGAL^®^ CA-630 [NP-40] 1%, sodium deoxycholate 0.5% and SDS 0.1%), supplemented with 1X Pierce™ protease inhibitor (Thermo Fisher Scientific) and 40 µM MG-132. Samples were mixed with 2 mL of extraction buffer per gram of tissue powder, incubated on ice for 20 min with occasional vortexing and centrifuged at 18,000 x g for 20 min at 4 ⁰C. The supernatant was transferred to a PD-10 desalting column (Cytiva Life Sciences) to remove the free biotin from the total protein extract, following the manufactureŕs instructions. Before adding the total extract, the PD-10 columns were equilibrated with 12 mL of RIPA buffer (without protease inhibitor and without MG-132), followed by 4 mL of RIPA buffer supplemented with 1X Pierce™ protease inhibitor and 40 µM MG-132. The volume of samples was adjusted to be a multiple of 2.5 mL, because each PD-10 column is intended for a sample volume up to 2.5 mL. Samples were eluted with 3.5 mL of RIPA buffer supplemented with protease inhibitor and 40 µM MG-132, and the eluates from the same samples were combined in one pre-cooled tube. Biotinylated proteins were subsequently pulled down using Pierce™ Streptavidin Magnetic beads (Thermo Fisher Scientific), following the manufacturer’s instructions. Briefly, 200 µL of streptavidin beads (50% slurry) were equilibrated with a modified RIPA buffer that does not contain NP-40 (25 mM Tris-HCl pH 7.6, 150 mM NaCl, 0.5% sodium deoxycholate and 0.1% SDS). For western blot (WB) analysis, only 50 µL of streptavidin beads (50% slurry) were used [57]. Protein extracts were incubated with conditioned beads for 1 h at 4°C in an overhead rotator. The beads were washed three times with the modified RIPA buffer without NP-40, containing 1X Pierce™ protease inhibitor. For PL experiments, the beads were additionally washed three times with 50 mM ammonium bicarbonate (ABC) buffer, at pH 8 (Sigma Aldrich) and resuspended in 45 µL of the same ABC buffer. Enriched proteins for WB or LC-MS analysis were stored at -20 ⁰C until further use. The desalting and pulldown procedures were performed in a 4 ⁰C room.

### Western blot analysis

Proteins captured by the streptavidin beads were eluted in 60 µL 4X Laemmli Sample Buffer (Bio-Rad) and incubated at 95 ⁰C for 10 min, at 250 rpm. The eluted proteins were separated on a 4-20 % mini-PROTEAN TGX pre-cast gel (Bio-Rad) and transferred to 0.2 µm PVDF membrane (Bio-Rad), using the Trans-Blot Turbo system at 25 V and 1.3 A, for 7 min. The membrane was then blocked with StartingBlock Blocking Buffer (Thermo Fisher Scientific), supplemented with 0.5% Tween-20, for 30 min at room temperature (RT) on a tube roller. Then, the gel blot was incubated with αGFP-HRP (1:1000 in StartingBlock buffer with 0.5%Tween-20; Thermo Fisher Scientific, #A10260), to detect the YFP-TbID fusion proteins, and streptavidin-HRP (1:5000 in StartingBlock buffer with 0.5%Tween-20; Thermo Fisher Scientific, #21124) to detect the biotinylated proteins, at 4⁰C overnight in a tube roller. Chemo-luminescent signal was revealed using the SuperSignal West Dura substrate (Thermo Fisher Scientific) and further visualized in a ChemiDoc MP imaging system and analyzed with Image Lab software (Bio-Rad).

### Pre-processing of samples for LC-MS/MS

Biotinylated proteins enriched by streptavidin pulldown and present on the beads, were resuspended in 45 µL of ABC buffer and processed as described before for LC-MS/MS [77].

### LC-MS/MS and MaxQuant protein identification

For the LC-MS/MS analysis, the peptides were separated by reverse-phase nano liquid chromatography using a Thermo nLC1000 column, after which the peptides were measured using an Orbitrap Exploris 480 mass spectrometer. The peptide spectra were searched in Maxquant (version 2.0.3.0) 130, using the Andromeda search engine 131 with label-free quantification (LFQ), against the version UP000011115 of the *S. tuberosum* cv DM1-3 516 R44 proteome dataset, including the protein sequence of NbSOBIR1 (A0A0H3U2A6; SGN: Niben101Scf03816g01001.1), ELR (A0A5B8YUL8), INF1 (Q01905), LTI6B (Q9ZNS6) and frequently occurring contaminants.

### MS data analysis in Perseus

To identify putative significant NbSOBIR1-interacting proteins, we performed a statistical analysis with the LFQ values quantified from MS with Perseus software (version 1.6.2.3) [78]. We calculated two-sample t-tests and performed a permutation-based false discovery rate (FDR) correction (q-value) based on LFQ values. We used an FDR value of 0.05 and an S0 of 0.1. We considered putative interactor proteins as significant when the q-value was less than 0.05. Three comparisons were performed: (1) NbSOBIR1 versus AtLTI6B; (2) NbSOBIR1 (+E/I) versus AtLTI6B and (3) NbSOBIR1 versus NbSOBIR1 (+E/I). To visualize the analysis of each comparison, the log2-fold change and -log2 p-value from two-sample t-tests were plotted. A higher numerical value for the log2-fold change indicates a larger difference in the abundance of the biotinylated protein between two samples, whereas a higher numerical value for -log2 p-value indicates a higher level of statistical significance, meaning that the difference in abundance is more likely to be a true difference rather than a random fluctuation. Taken together, a higher value for both log2-fold change and -log2 p-value suggests a greater enrichment of a particular biotinylated protein under the experimental condition. KEGG pathway and GO Molecular Function analysis were performed with the ShinyGO 0.77 webtool [79], using a FDR cutoff of 0.05. KEGG pathways with a p-value ≤ 0.05 and GO Molecular Function with a p-value ≤ 0.001 were plotted. The prediction of cellular localization was based on the subcellular localization and GO Cellular Components information that UniProt provides for the identified proteins. BLAST of selected proteins was performed on UniProt against *Solanum lycopersicum* (Tomato) (ID: UP000004994 and *Arabidopsis thaliana* (cv. Columbia) (ID: UP000006548).

## Author Contributions

T.M.F., M.H.A.J.J. and V.G.A.A.V. conceived this study. T.M.F., S.L.V., M.H.A.J.J., and V.G.A.A.V. designed experiments and T.M.F. and S.B., performed them. T.M.F., S.L.V., S.B., and M.H.A.J.J. analyzed and/or interpreted data. T.M.F. wrote the paper, with contributions from M.H.A.J.J. and V.G.A.A.V.. All authors contributed to finalizing the manuscript and approved the final version.

## Acknowledgments

This work was supported by European Union’s Horizon 2020 research and innovation program under grant agreement no. 766048 (MSCA-ITN-2017 PROTECTA), and the Peruvian National Council for Science, Technology and Technological Innovation (CONCYTEC) and its executive unit FONDECYT. We thank Li Shi for her help during the proximity-labeling assays and useful discussions.

## Supplementary information

**Figure S1.** Constructs and encoded protein sequences used for proximity-dependent labeling experiments.

**Figure S2.** Screening of Solanum genotypes with a genetic background of *S. microdontum*.

**Table S1.** Label free quantification (LFQ) values and ratio of YFP, TurboID and NbSOBIR1 proteins per treatment (AtLTI6B-YFP-TbID, NbSOBIR1-YFP-TbID and NbSOBIR1-YFP-TbID +(E/I)).

**Table S2.** Identification of proteins that interact with full SOBIR1 or SOBIR1 + E/I using TurboID-based proximity labeling.

## Notes

### Competing Interest Statement

The authors have declared no competing interest.

