## Supplementary material for "TurboID-based proximity-dependent labeling using SOBIR1 as a bait in potato leads to the identification of novel defense-related signaling partners": Figure S1

**A**

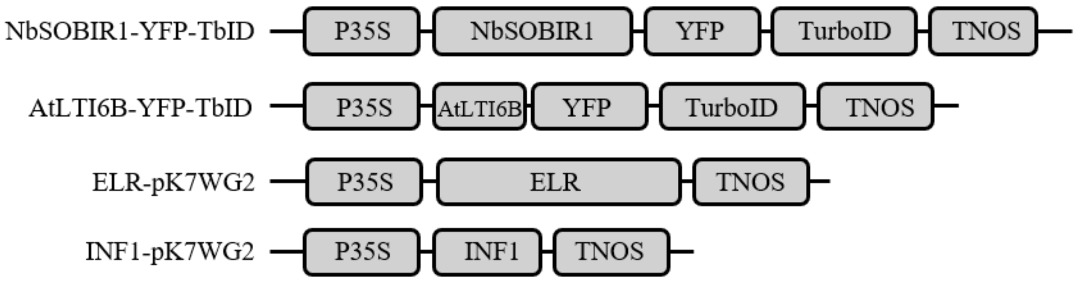


**B**

**NbSOBIR1 (A0A0H3U2A6)**

MAFTASQIHFFFFSLFAFLLIAVQARLNLYPPDHAALLLVQKDLGIQGQRIALCNSATISCERRKANRTQLLRVTRIDFRSSGLSGT
LSPAIGKLSVLKELSLPNNQLFDQIPVQILDCRKLEILDLGNNLFSGKVPSELSSLLRLRILDLSSNEFSGNLNFLKYFPNLEKLSLA
DNMFTGKIPPSLKSFRNLRILNISGNSFLEGHVPVMSQVEHLSAELDQHFVPKRYILAENSTRSNQISALAPNSNSGNAPAPAPSH
NVTPIHKHSNRKKRKVRAWLLGFFAGSFAGAISAVLLSVLFKLVMFFVRKGKTDGTLTIYSPLIKKAEDLAFLETEDGVASLEMI
GKGGCGEVYRAELPGSNGKIIAIKKIIQPPMDAAELTEEDTKALNKKMRQVKSEIQILGQIRHRNLLPLLAHMPRPDCHYLVYEY
MKNGSLQDILQQVTEGTRELDWLGRHRIAVGIASGLEYLHINHSQCIIHRDLKPANVLLDDDMEARIADFGLAKALPDAHTHVT
TSNVAGTVGYIAPEYHQTLKFTGKCDIYSFGVVLAVLVIGKLPSDEFFQHTPEMSLVKWLRNVMTSEDPKRAIDSKLIGNGFEEQ
MLLVLKIACFCTLENPKERPNSKDVRCMLTQIKH

**AtLTI6B (Q9ZNS6)**

MSTATFVEIILAIILPPLGVFLKFGCKVEFWICLILTLFGYLPGILYALYIITK

**ELR (A0A5B8YUL8)**

MVMSLFFFYSFLCFVFLISGCFSSSFDHHLCSPTEASSLLQFKQSFQISDYSTLKCDTSFPKTKSWNESRDCCSWDGVTCDLLNGH
VIGLDLSCSQLRGSIHPNSSLFQLHHLQTVNLAYNNFSTSSISHNIGRWRNLRHLNLSNSFFSGKIPTEISFLSNLVSLDLSSSYGLQ
LDERTFETMLHNFTNLEVLALFLGNISSPIPVSIHPNSSLFQLHHLHTLNLVNNFFYPSSIPNGIGRLRNLRHLKLYGFQGKIPTEISY
LSNLVSLDLSYSYERLQLDERTFEAMFQNLTNLELLSLYGVNISSQIPVNISSSLRYLDLGYTNLRGVLTENFFLLPNLEILKLSGN
DLVKGVFPKIHWSNTLLMELDISSTGISGEVPDSIGTFSSLNILNLAGCQFSGSIPDSISNLTQIRELILYHNHFTGHIPSTISKLKHLT
RLDLSINYFSGEIPDVFSNLQELRTLHLSYNSFIGSFPASILSLTHLEYLGLSSNSLSGPLPSNQSMLQKLTELNLSYNSLNGTIPSWV
FSLPLLSSVSLQHNRLRGLADEVIKTNPTLKQLYLSNNQLSGSFPQSLVNLTNLETLGISSNNITIDEGMNITFLSLSSLFLSSCQLK
DFPHFLRNIKTLRYLDISNNKICGQIPNWFSGMRWDSLQFLNLSHNSLTGHLPQFRYDNLQYLDLKFNYLQGPLALFICNMRKLI
LLDLSHNYFSDSVPHCLGSMPNLRVLDLRRNNFTGSLPPLCAQSTSLSTIVVNGNRFEGPVPVSLLNCNGLQVLDVGNNAINDTF
PAWLGTLQELQVLILKSNKFHGPISTCHTKFCFPKLRIFDLSRNDFSGSLPAKVFRNFKAMIKLDGEDTGNIKYMESMSNLPFVRS
YEDSVSLVIKGQDIELQRISTIMTTIDLSNNHFEGVIPKTLKDLSSLWLLNLSHNNLLGHIPMELGQLNTLEALDLSWNWLTGKIPQ
ELTRMNFLAVLNLSQNHLVGPIPQGPQFNTFENDSYCGNLDLCGPPLSKQCETSDSSHVPQQLESEEEGESYFFSGFTWESVVIGY
SFGLVVGTVMWSLMFKYRKPKWFVEFFDGLMPHKRRRPKKRAQRRRT

**INF1 (Q01905)**

MNFRALFAATVAALVGSTSATTCTTSQQTVAYVALVSILSDTSFNQCSTDSGYSMLTATSLPTTEQYKLMCASTACKTMINKIVS
LNAPDCELTVPTSGLVLNVYSYANGFSSTCASL

Figure S1. Constructs and encoded protein sequences used for proximity-dependent labeling experiments.
