## Supplementary material for "TurboID-based proximity-dependent labeling using SOBIR1 as a bait in potato leads to the identification of novel defense-related signaling partners": Figure S2

| Genotype | INF1 | ELR+ INF1 | EV |
|---|---|---|---|
| GIG362-6 | 0 | 0 | 0 |
| MCD360-1 | 0 | 0 | 0 |
| 3521-321 | 0 | 10 | 0 |
| 3521-169 | 9 | 10 | 0 |
| 3521-1698 | 8 | 10 | 0 |
| 3521-172 | 0 | 10 | 0 |
| 3521-183 | 0 | 10 | 0 |
| 3521-460 | 0 | 10 | 0 |
| 3521-589 | 5 | 10 | 0 |
| 3521-80 | 8 | 10 | 0 |
| 3521-43 | 0 | 10 | 0 |
| 3521-59 | 8 | 10 | 0 |
| 3521-849 | 7 | 6 | 0 |
| 3521-18 | 0 | 6 | 0 |
| 3521-1153 | 0 | 0 | 0 |
| 3521-1853 | 0 | 0 | 0 |
| 3521-1856 | 4 | 0 | 0 |
| 3521-1581 | 3 | 10 | 3 |
| 3521-1672 | 8 | 10 | 3 |
| 3521-307 | 7 | 10 | 3 |
| 3521-1222 | 5 | 9 | 5 |
| 3521-935 | 0 | 10 | 8 |
| 3521-28 | 10 | 10 | 8 |
| 3521-1221 | 10 | 10 | 10 |
| 3521-1635 | 10 | 10 | 10 |
| 3521-353 | 10 | 10 | 10 |
| 3521-388 | 10 | 10 | 10 |

**Figure S2.** Screening of Solanum genotypes with a genetic background of *S. microdontum*.
