## Supplementary material for "TurboID-based proximity-dependent labeling using SOBIR1 as a bait in potato leads to the identification of novel defense-related signaling partners": Table S1

**Table S1. Label free quantification (LFQ) values and ratio of YFP, TurboID and NbSOBIR1 proteins per treatment (AtLTI6B-YFP-TbID, NbSOBIR1-YFP-TbID and NbSOBIR1-YFP-TbID +(E/I)).**

| | Sample | | | Ratio LFQ | | |
|---|---|---|---|---|---|---|
| **Protein ID** | AtLTI6B | NbSOBIR1 | NbSOBIR1 (+E/I) | AtLTI6B/ NbSOBIR1 | NbSOBIR1/ NbSOBIR1(+E/I) | AtLTI6B/ NbSOBIR1(+E/I) |
| **YFP** | 3,53E+08 | 1,06E+08 | 3,64E+07 | 3,33 | 2,91 | 9,70 |
| **TurboID** | 5,77E+09 | 2,08E+09 | 6,27E+08 | 2,77 | 3,32 | 9,20 |
| **NbSOBIR1** | 4,67E+05 | 1,09E+09 | 3,16E+08 | 4,28E-04 | 3,45 | 1,48E-03 |
